# Pannexin-2 deficiency disrupts visual pathways and leads to ocular defects in zebrafish

**DOI:** 10.1101/2024.10.01.616190

**Authors:** Riya Shanbhag, Georg S.O. Zoidl, Fatema Nakhuda, Shiva Sabour, Heike Naumann, Christiane Zoidl, Armin Bahl, Nima Tabatabaei, Georg R. Zoidl

## Abstract

Pannexin-2 (Panx2) is a unique ion channel localized to ER-mitochondria contact sites. These dynamic and specialized microdomains are abundant in neurons and glia and essential for cellular signaling and metabolism. While synaptic interactions are well-studied, the role of intracellular contacts, such as those of ER-mitochondrial junctions, in neuronal function and neurodegeneration remains largely unexplored. To investigate the roles of Panx2 in neuronal communication, we meticulously examined its expression pattern in the zebrafish brain and used TALEN technology to generate homozygous Panx2 knockout (*Panx2^Δ11^*) zebrafish. Our results demonstrate that *panx2* mRNA is present in several brain regions, notably in visual centers such as the optic tectum and the thalamus. In 6 days post fertilization TL (*Panx2^+/+^*) larvae, Panx2 expression was observed in the inner and outer plexiform layers of the retina and the arborization fields of the optic tract. Transcriptome profiling of *Panx2^Δ11^* larvae by RNA-seq analysis revealed down-regulation of vision-related genes, specifically those involved in visual and sensory perception and lens development. Behavioral tests showed that loss of Panx2 leads to an altered ability to interpret visual information, such as changes in ambient illuminations, and respond with the characteristic motor action. *Panx2^Δ11^* larvae exhibited reduced locomotor activity during light and increased activity during dark phases. Additionally, the knockout larvae displayed significantly impaired optomotor response (OMR). Lastly, when we tested the retinal structure of adult zebrafish eyes using optical coherence tomography (OCT), *Panx2^Δ11^* fish revealed a longer mean axial length and a negative shift in retinal refractive error (RRE) values; both indicative of myopia. Our findings highlight a distinct, novel function of Panx2 in retinal development, visual perception, and ocular health, beyond its recognized roles in neurodevelopment and tumor-suppressing properties in cancer cells.

## Introduction

Pannexin-2, the least studied member of the vertebrate-specific family of three proteins: Pannexin-1 (Panx1), Pannexin-2 (Panx2), and Pannexin-3 (Panx3) ^1^, has been shown to exhibit a dynamic expression pattern. Early reports demonstrated low levels of mammalian Panx2 during prenatal development, with a significant increase postnatally, suggesting its involvement in CNS expansion ^2^. This finding, along with another early report pointing toward the role of Panx2 in regulating the timing of neuronal differentiation via an S-palmitoylated modification ^3^, underscores the significant implications of Panx2’s role in the central nervous system. Although originally believed to be confined to the central nervous system ^4–6^, Panx2 has since been detected in sixteen different tissues; highest in skin, skeletal muscles, and the eye ^7^. Recent structural studies have revealed Panx2 as a channel-forming protein with a unique pore architecture that allows the passage of small molecules, including ATP ^8,9^. This unique structure, along with its post-translational modifications, including N-glycosylation at asparagine (N86) within its first extracellular loop ^10^, sets Panx2 apart from its paralogs, Panx1 and Panx3. Unlike Panx1 and Panx3, N-glycosylated Panx2 is absent from the plasma membrane ^11,12^. While Panx2 was known to localize within the cytoplasm, its specific organelle distribution was not consistently defined. Studies reported its presence in various intracellular structures, including the ER, Golgi apparatus, and endolysosomes ^10,13,14^. Le Vasseur et al. identified Panx2 as a novel component of mitochondria-associated membranes (MAMs), using compelling live-cell imaging and electron microscopy ^15^. The authors showed that approximately 60% of endogenous Panx2 is localized at the mitochondria under physiological conditions, suggesting specialized roles that differentiate it from other pannexins. MAMs are dynamic ER-mitochondria contact sites crucial for cellular processes such as lipid transfer, calcium homeostasis, immune regulation, and cell death ^16^. We know there are numerous ER-mitochondria contact sites in neurons ^17^. However, the specific roles of Panx2 within these junctions, particularly in the context of neurotransmission, neurodegenerative diseases, and cancer, remain to be elucidated ^18,19^.

Multiple Panx2 isoforms exist in humans and zebrafish, with conserved regions primarily localized within the transmembrane domains and intracellular loops ^6^. Notably, the Panx2 C-terminal domain exhibits significant sequence variation compared to Panx1 and Panx3 ^9^. To investigate whether these paralog differences correlate with distinct expression patterns, we performed HCR-fluorescent *in situ* hybridization of whole larval zebrafish. Panx2 mRNA was detected in multiple brain regions. To further elucidate the role of Panx2 in neuronal communication, we generated a global Panx2 knockout zebrafish model using TALEN technology. A custom-made antibody identified Panx2 in the inner segments of photoreceptors and the outer retina and confirmed the loss of Panx2 in the knockout. Transcriptome analysis revealed differential expression of genes involved in visual perception, circadian rhythm, and innate and adaptive immune responses. Subsequent behavioral studies demonstrated that loss of Panx2 impairs visual-motor responses and optomotor behavior, highlighting its critical role in visual information processing. Lastly, optical coherence tomography of adult zebrafish eyes revealed alterations in the structure of the lens, suggesting that early-life molecular changes induced by the absence of Panx2 result in a myopia-like phenotype. We concluded that Panx2’s novel association with myopia contributes to the observed visual-motor phenotype.

Our study reveals a novel and essential role for Panx2 in the visual system. Given the expression patterns of Panx2 in the brain and its subcellular localization at mitochondria-associated membranes (MAMs), it is likely that this protein plays diverse roles in neuronal function. While Panx2 has been implicated in various processes, including skin homeostasis, neurodevelopment, and cancer, our findings expand its known functions to include visual perception. This discovery highlights the multifaceted nature of Panx2 and its potential significance in neurological disorders.

## Results

### Panx2 is expressed broadly in the brain of zebrafish larvae

RNA fluorescence in-situ hybridization (RNA-FISH) was used to determine *panx2* mRNA expression in 6 days post-fertilization (dpf) whole-mount TL larvae. By analyzing volumes in the common reference frame of the mapZebrain altas ^20^, we found *panx2* distributed within the prosencephalon, mesencephalon, and rhombencephalon (**Fig. 1a,b**). *Panx2* mRNA levels were highest in the mesencephalon (median fluorescence intensity/pixel for *panx2* probe = 1191, n=6; control = 425, n=5; P-value <0.0001) (**Fig. 1c**). In light of our prior reports on the roles of Panx1a/b in zebrafish visual function, we focused on brain regions involved in the ascending visual pathway (**Fig. 1d,e**). We found high *panx2* localized within the pallium, pretectum, tectum neuropil, tegmentum, periventricular layer (PVL), and dorsal thalamus (**Fig. 1f,g**). The elevated *panx2* expression in the neuropil, a region crucial for visual integration, suggested a potential involvement of Panx2 channels in synaptic plasticity and neuronal communication within this visual processing hub (median fluorescence intensity/pixel for *panx2* probe = 2570, n=6; control = 750, n=5; P-value <0.0001). Moreover, *panx2* was concentrated in arborization fields (AFs) 4, 6, 7, 8, and 9 (**Fig. 1h,i**). Previous studies have shown that these AFs are key players in the zebrafish visual system. Specifically, AF4 is involved in phototaxis; AF8 and AF9 are important for the response to dimming and brightening environments ^21^. While AF6 has a potential role in optokinetic and optomotor reflexes. Quantification of *panx2* fluorescence within the AFs revealed highest levels in AF7 and AF9, both of which are associated with the pretectum (**Fig. 1j**).

**Figure 1:**
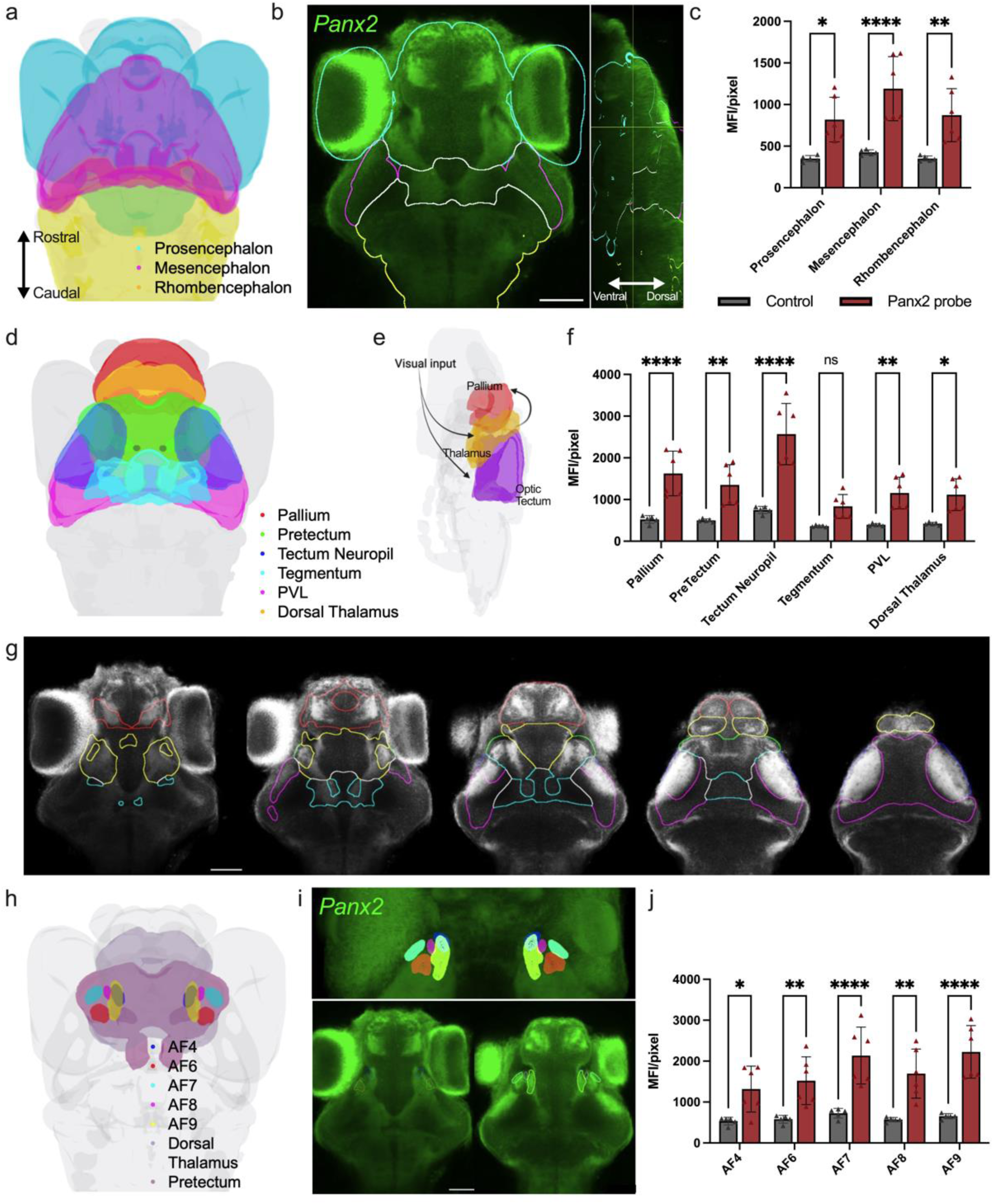
Spatial distribution of *panx2* mRNA by fluorescence in-situ hybridization. a) Three-brain region atlas derived from Kunst et al. (2019) showing the major brain subdivisions: prosencephalon, mesencephalon, and rhombencephalon. b) Dorsal view of TL larvae expressing *panx2* (green). Scale bar: 50 µm. c) Median fluorescence intensity (MFI) for regions described in (a), panx2 probe (red), control (same procedure without probe, grey). d) Three-dimensional view of larval zebrafish brain regions involved in the visual ascending pathway, acquired from mapZebrain. e) Simplified schematic of the visual processing pathway; visual input travels to the thalamus and optic tectum, subsequently projecting to the pallium. f) MFI for regions described in (d). g) Images from left to right represent sequential slices of *panx2*-labelled larval brain regions involved in the ascending visual pathway: axial views, ventral to dorsal. h) Three-dimensional view of larval zebrafish brain regions showing retinal arborization fields (AFs) 4, 6, 7, 8 and 9 that partially project within the dorsal thalamus and pretectum. i) Three-dimensional projection showing AFs mentioned in (h) (top panel). Images of *panx2* expression within selected AF regions (bottom panel). j) MFI for regions described in (h). Scale bar: 50 µm. Statistical significance determined with two-way ANOVA. Significance: ****P-value<0.0001, ***P-value<0.001, P-value<0.01 and *P-value<0.05. Error bars = SD. Sample sizes for TL labelled with *panx2* probe, n=6; control probe, n=5.

### Generating a Panx2 global knockout zebrafish line

The *panx2* mutant allele was generated by TALEN-mediated genome editing, targeting the *BamHI* restriction endonuclease site in the first exon of the *panx2 gene* (**Fig. 2a**). A DNA sequence analysis of microinjected embryos identified short 4 to 11 base pairs (bp) long nucleotide deletions in exon one of the *panx2* gene. An adult founder fish (F0) with an 11 base pair deletion (*panx2*Δ11) was selected for further experimentation (**Fig. 2b**). In this fish the 11 bp deletion caused a frameshift at amino acid (aa) G28, resulting in a premature stop codon. The consequence of this stop codon was a truncated reading frame for a 27-aa protein, lacking 640-aa of the 651-aa long Panx2-C protein sequence (**Fig. 2c**). The *Panx2^Δ11^* fish were viable and able to breed for multiple generations. When comparing age-matched male and female adults to wild type siblings a modest but significant decrease in body length was noted (**Fig. 2d,e**). The gene-editing event in exon1 effectively eliminated the shared start codon for the three known Panx2 protein isoforms A: 580-aa (ABY78016), B: 604-aa (ABY78018), and C: 651aa (ABY78017). The Panx2-C isoform was the most abundant *panx2* mRNA detectable in 6 dpf TL (*Panx2^+/+^*) larvae by RT-qPCR suggesting that it may play a dominant role in Panx2-mediated functions (**Fig. 2f**).

**Figure 2:**
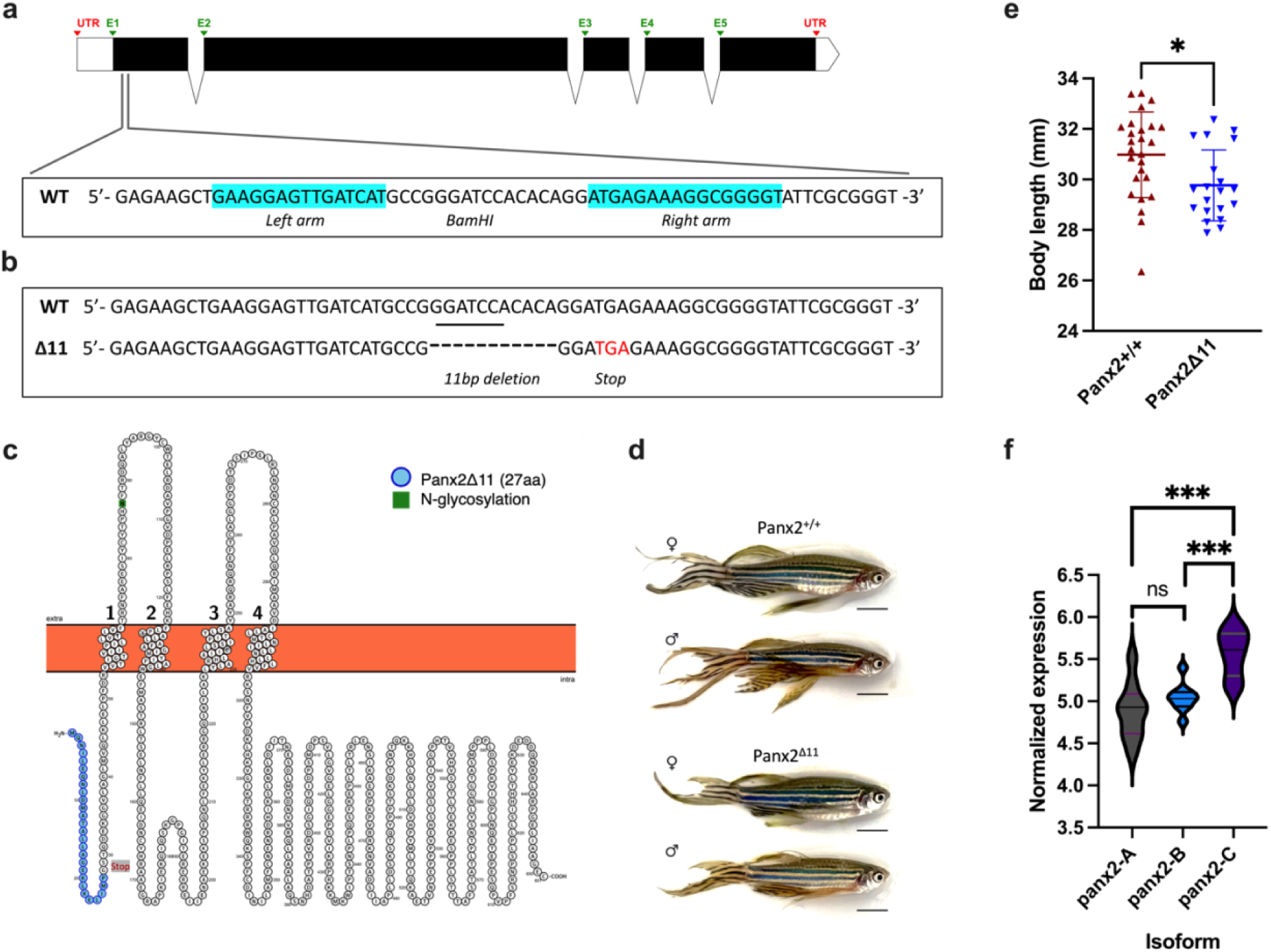
Generating a Panx2 global knockout zebrafish line. a) Exon-intron organization of the zebrafish Panx2 gene (ENSDARG00000063019, chromosome 18 (GRCz11:CM002902.2). The positions of the targeting Left and Right arms with a unique BamHI restriction endonuclease site in a spacer region between the TALEN pair are highlighted. b) Representation of the 11 bp deletion causing a premature stop codon. c) Graphical representation of the predicted Panx2 structure. The region in blue indicates the residual sequence of 27 amino acids followed by a new in-frame stop codon. Representative images (d) and comparison of body length between one-year-old zebrafish of different genotypes, both sexes. f) Real-time PCR quantification and presentation of the normalized expression of the three Panx2 isoforms, normalized to 18S rRNA. Statistics in e,f: Welch’s test. Sample sizes were n=26 for *Panx2^+/+^*and n=20 for *Panx2^Δ11^*. Significance: ***P-value<0.001 and *P-value<0.05. Error bars = SD.

### Panx2 is localized in the outer retina of the zebrafish

To demonstrate the protein-level knockout of Panx2 in larvae, a custom polyclonal anti-peptide Panx2 antibody was generated. *Panx2^+/+^*larvae revealed low Panx2 immunoreactivity, which was most pronounced in the outer retina, the inner plexiform layer, and the lens epithelium (**Fig. 3a, see open triangles**). Panx2 knockout larvae exhibited a significant reduction in fluorescence with residual signals attributed to collagen autofluorescence in the sclera and at the margins of the lens epithelium, confirming the loss of function at the translational level (**Fig. 3b**). At higher magnification of the outer retina Panx2 localization was prominent in the axons of the inner segments of photoreceptor cells and the lens epithelium (**Fig. 3c,d, see open triangles**). No immunoreactivity was detected in horizontal cells which express two other pannexins, the Panx1a and Panx1b isoforms ^22^.

**Figure 3:**
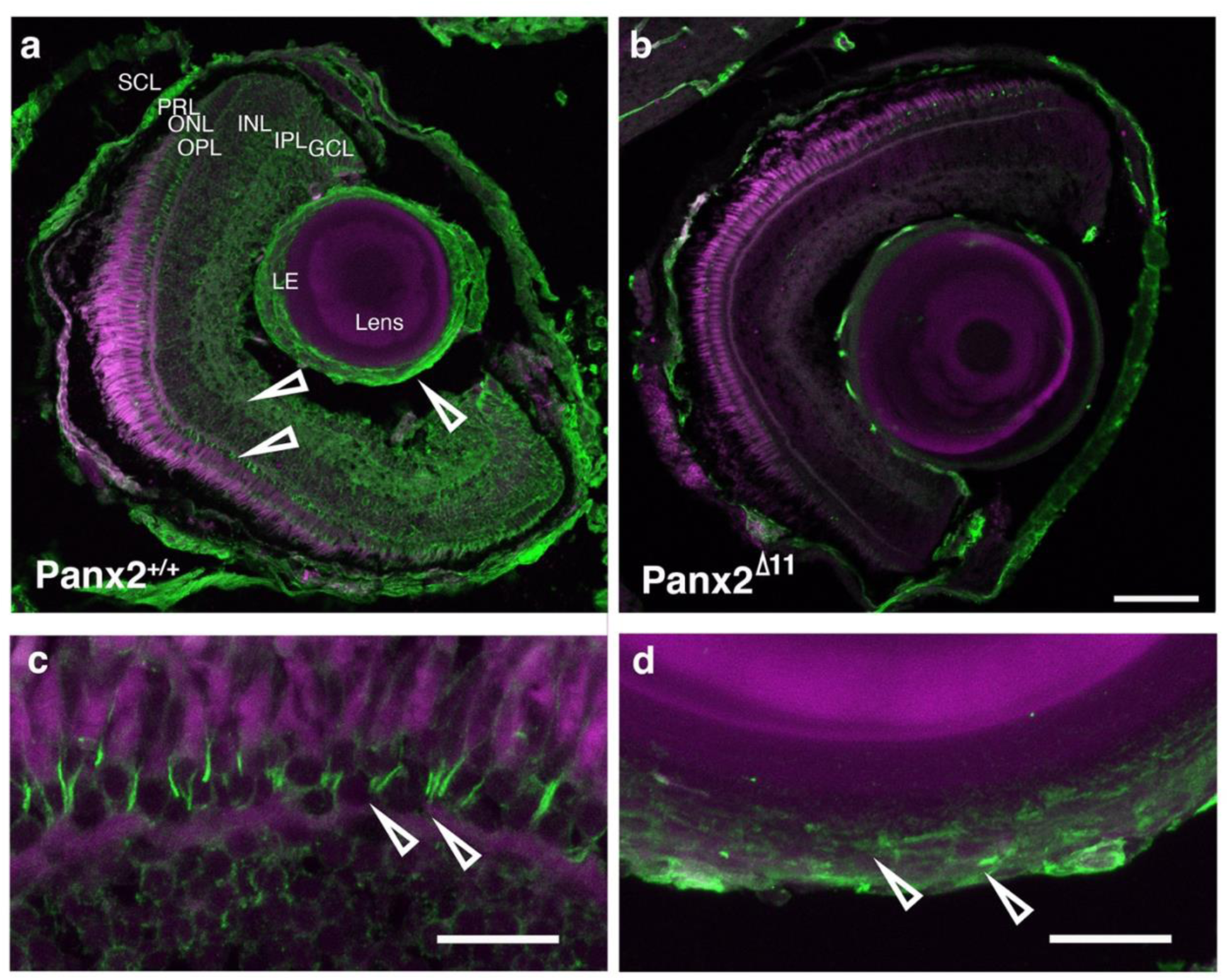
Panx2 localization in outer retina of the zebrafish. a) TL larvae showing that Panx2 proteins (green) are ubiquitously expressed throughout the retina. The expression was enriched in the outer retina, the inner plexiform layer, and the lens epithelium. DAPI is shown in magenta. b) After Panx2 ablation the expression was greatly reduced across the retina, with residual staining left in tissue known for high autofluorescence. c) Higher magnification of the outer retina showing the pronounced staining in sections of the inner photoreceptor segments, in TL larvae. d) Higher magnification of the lens epithelium of TL larvae. Abbreviations: SCL sclera, LE lens epithelium, GCL ganglion cell layer, IPL inner plexiform layer, INL inner nuclear layer, OPL outer plexiform layer, ONL outer nuclear layer, PRL photoreceptor layer. Scale bars: (a, b) 60 µm; (c, d) 10 µm.

### Panx2 ablation regulates vision processes and structural components of the lens

A comparison of the transcriptomes of 6 dpf *Panx2^+/+^* and *Panx2^Δ11^* larvae identified 5355 differentially expressed genes (DEG) when the adjusted P-value (padj) was <0.05. In total, 2512 genes were up-regulated, and 2843 were down-regulated (**Fig. 4a**). The Gene Ontology analyzer for - (GOSeq 1.56.0; ^23^) identified Biological Processes (BP) and Molecular Functions (MF) that were over/under-represented in the DEG data (**Fig. 4b**). Vision-related categories like *Sensory Perception of Light Stimulus (GO:0050953; 61 genes down)* or *Sensory Perception* (GO:0007600; 70 genes down) were significantly down-regulated (**Fig. 4c,d**). The GO annotations refer to the events required for an organism to receive a sensory light stimulus, convert it to a molecular signal, and recognize as well as characterize it. The most up-regulated process was the *Humoral Immune Response* representing immune responses mediated through a body fluid *(GO:0006959; 17 genes up)* (**Fig. 4e**)*. The Structural Component of the Lens (GO:0005212; 38 genes down)* was the most significantly down-regulated Molecular Function. This GO annotation refers to molecules that contribute to the structural integrity of the lens (**Fig. 4f**). The most up-regulated MFs were *Endopeptidase Regulator Activity (GO:0061135)* encompassing molecules that modulate the activity of peptidases, which can control critical functions of innate and adaptive immune responses like antigen processing and presentation of immunogenic peptides.

**Figure 4:**
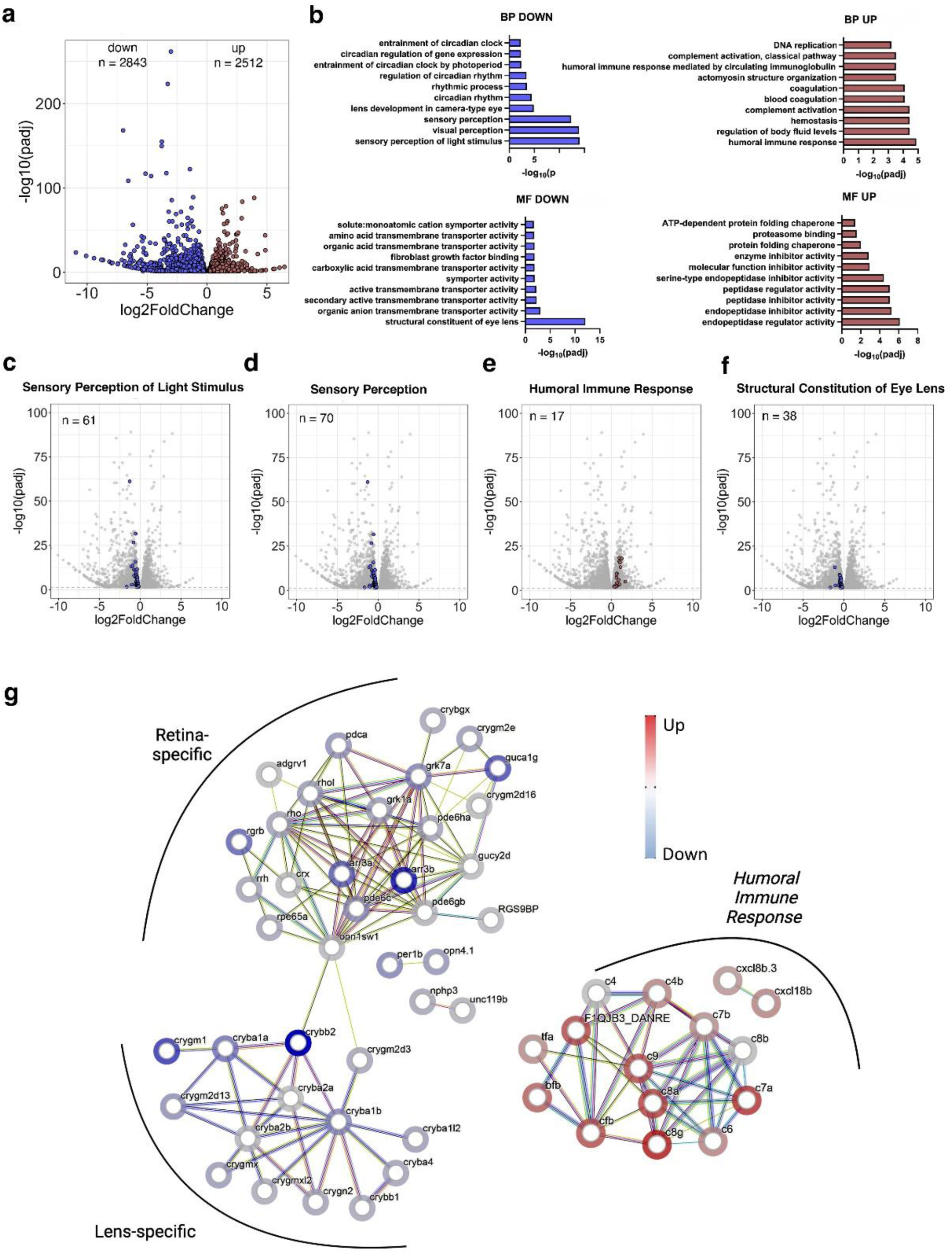
Panx2 ablation regulates vision processes and structural components of the lens. a) A DESeq2 analysis of RNA-seq data from Panx2^+/+^ and Panx2^Δ11^ 6 dpf zebrafish larvae identified 5355 differentially expressed genes with an adjusted P-value < 0.05 b) Analysis of differentially expressed genes using GOSeq (v1.56.0). The Gene Ontology Classifier Biological Process (BP) showed significant downregulation of several vision-related categories. c,d) The most significantly down-regulated biological processes included *Sensory Perception of Light Stimulus (GO:0050953)* and *Sensory Perception (GO:0007600).* e) The top up-regulated BP was *Humoral Immune Response (GO:0006959).* f) The most significantly down-regulated Molecular Function (MF) was *the Structural Component of the Lens (GO:0005212).* g) STRING analysis of 61 regulated genes annotated as *Sensory Perception of Light Stimulus (GO:0050953)* and 17 regulated genes annotated as *Humoral Immune Response (GO:0006959).* K-means clustering identified two clusters which were either retina- or lens-specific. Cluster I genes are implicated in the phototransduction processes of the retina. Cluster II represents lens-specific proteins belonging to the alpha- or beta/gamma supergene families. A third group represents the humoral immune response.

Next, a STRING analysis with k-means clustering of the 61 differentially down-regulated genes annotated as *Sensory Perception of Light Stimulus (GO:0050953)* showed that both the retina and the lens were affected by loss of Panx2. Cluster I represented genes with essential roles in normal vision and signal transduction in the retina. Genes belonging to the retinal arrestin family (*arr3a* and *arr3b*), rhodopsin family (*rho* and *rhol*), as well as the guanylyl cyclase family (*guca1g* and *gucy2d*), were significantly down-regulated (**Fig. 4g**). Cluster II represented lens-specific proteins belonging to the two alpha- or beta/gamma groups of the crystallin super gene families, including members of the gamma-N and gamma-M subgroups. The group represented the M-subfamily of gamma-crystallin, which is specific to fish. These genes play essential roles in the lens structure; mutations have been shown to cause cataracts.

Genes in the up-regulated category *Humoral Immune Response (GO:0006959)* (**Fig. 4g**) showed notable up-regulated genes in *Panx2^Δ11^*larvae. Genes such as *si:dkey-22f5.9, c8a, c8g, c9*, are expressed in the liver. Interestingly, genes implicated in human eye diseases (*cfb*) and age-related macular degeneration (c2/*si:ch1073-280e3.1 (F1QJB3_DANRE*) were also found to be up-regulated. The significant differential expression of retina and lens genes, as well as genes implicated in human eye disorders, led us to prioritize biological processes related to vision.

### Light-ON and light-OFF conditions affect free swimming of Panx2^Δ11^ larvae

We examined visually guided behaviors to explore consequences of the differential expression of genes with roles in visual function. Baseline spontaneous swimming activity of both genotypes was analyzed at 6 dpf, during light-ON (1200 lux) and light-OFF (0 lux) phases. Larvae were placed individually in wells of a 24-well plate (each well: 1.9 cm^2^). The representative images illustrate locomotor patterns within a single well in the light-ON condition (**Fig. 5a**). Knockout larvae preferred swimming close to the circumference of the well. In the light, *Panx2^Δ11^* larvae swam a shorter distance (t=7.755, df=113.1, P-value <0.001; n=24 larvae) at a lower velocity (t=7.177, df=117.6, P-value <0.001; n=24 larvae) than the *Panx2^+/+^*wildtype group (**Fig. 5b,c**). In the absence of light, both *Panx2^+/+^* and *Panx2^Δ11^* larvae exhibited a preference for the perimeter and avoided the central zone of the well (**Fig. 5d**). However, in the light-OFF condition, *Panx2^Δ11^* larvae swam further (t=4.147, df=88.91, P-value <0.001; n=24 larvae) at a lower, steady velocity than *Panx2^+/+^* larvae (t=4.095, df=96.20, P-value <0.001; n=24 larvae) (**Fig. 5e,f**).

**Figure 5:**
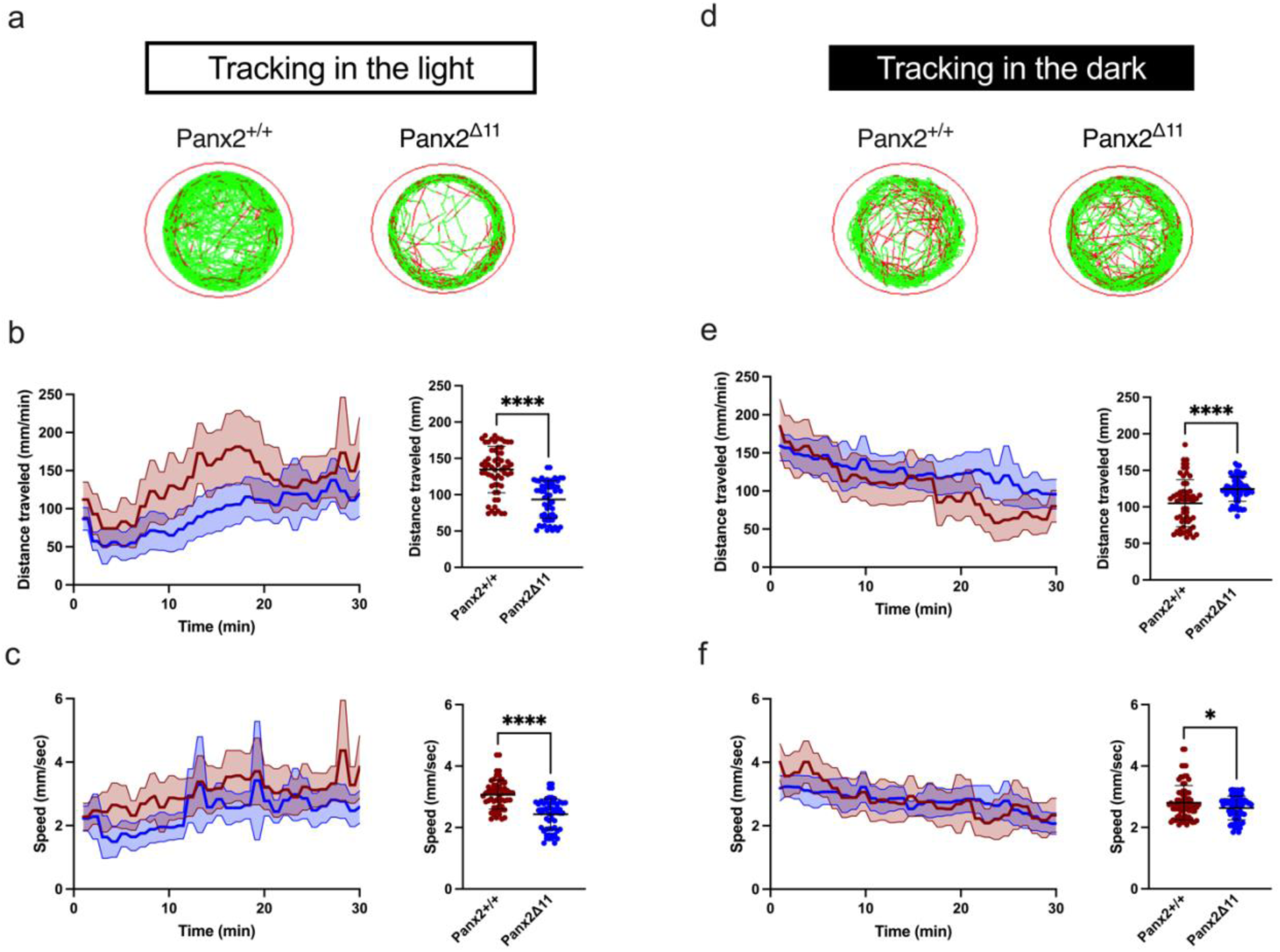
Light-ON and light-OFF conditions affect free swimming of *Panx2^Δ11^*. Representative images of a single well showing examples of *Panx2^+/+^* and *Panx2^Δ11^* larvae locomotion patterns in the light (a). Medium (<20 mm/sec) or high-speed movements (>20 mm/sec) are visualized with green and red colors, respectively. Graphs demonstrate mean experimental traces ± 95% CI of distance traveled (mm) and speed (mm/sec) of 6 dpf larvae for 30 min (b,c) and their corresponding mean values, in the light. d) Representative images for each genotype, in the dark. Tracked experimental traces of distance and speed under dark conditions (e,f) and their corresponding mean ± SD. Sample sizes were n=24 larvae for each genotype. A Welch’s t-test was used for statistical comparison between groups. Significance: ****P-value<0.0001 and *P-value<0.05. Error bars = SD.

### Panx2 ablation affects the visual motor response to light-ON and light-OFF stimuli

Visual motor response (VMR) analysis was performed to compare the stereotypical motor responses of *Panx2^+/+^* and *Panx2^Δ11^*larvae. The kinematic data collected provided information about seven parameters: duration of bouts, duration of bursts, freeze duration, average activity duration, bout counts, burst counts, and freeze counts. A principal component analysis identified “freeze duration” (dimension 1) and “average activity duration” (dimension 2) as capturing 73.62% and 15.14% of the data variance, respectively (**Fig. 6a**). Both parameters were selected to compare divergence between the genotypes. We found that *Panx2^Δ11^*larvae displayed strikingly different activity patterns for freeze duration, which represents the amount of time that the larvae were immobile (**Fig. 6b**) and average activity duration (**Fig. 6c**). Panx2 knockout larvae were less active during light-ON phases, and more active in light-OFF phases. A Welch’s t-test established statistical differences in mean activity of *Panx2^+/+^*and *Panx2^Δ11^* in both luminance conditions (P-value <0.0001 for light-ON (**Fig. 6d**), P-value <0.0001 for light-OFF (**Fig. 6e**), n=24 larvae).

**Figure 6:**
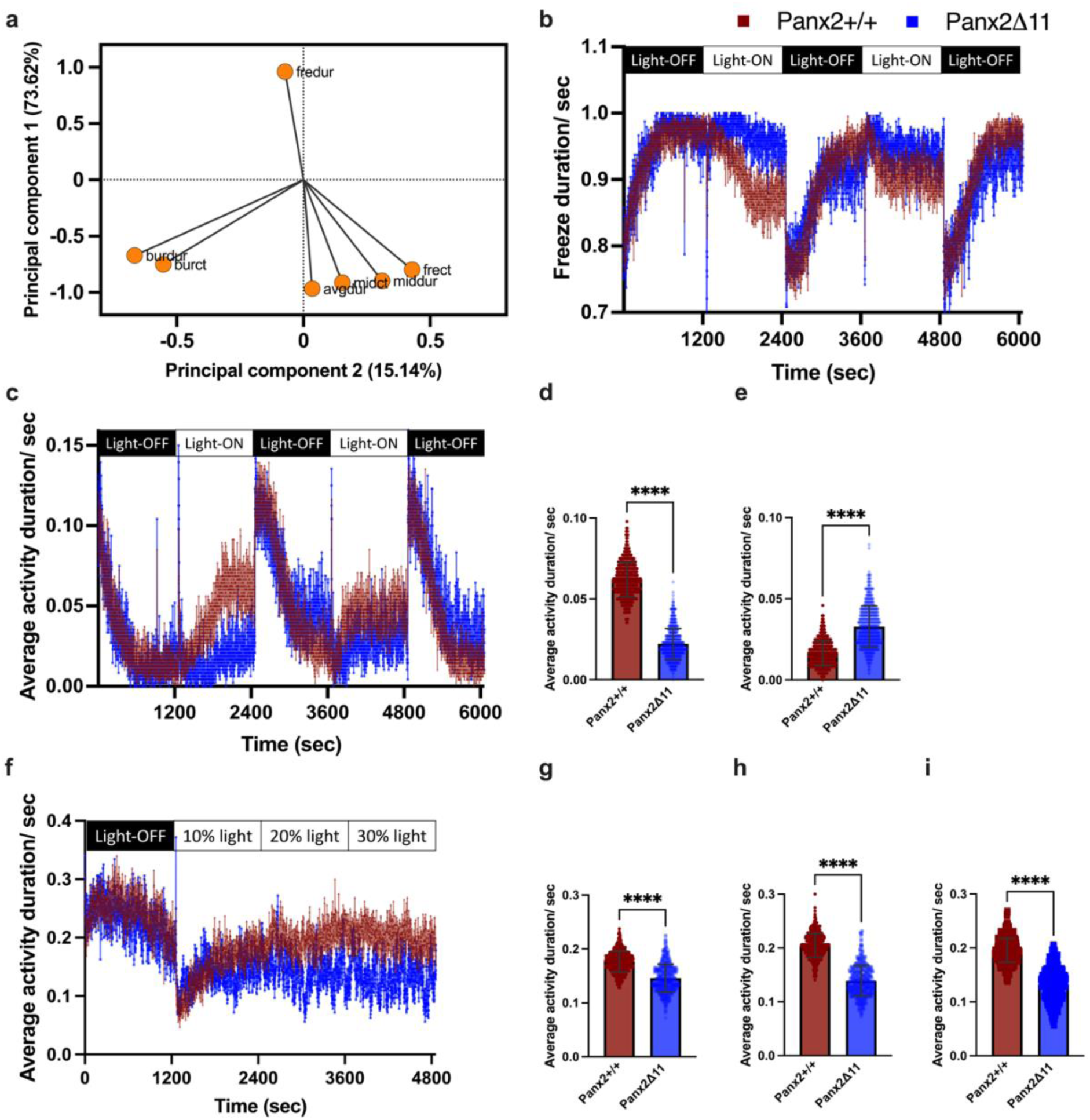
Panx2 ablation affects the response to light-ON and light-OFF stimuli. Principal component analysis was performed using multivariable activity data. a) The variable correlation plot indicates the coordinates of seven variables within the first two dimensions, where PC1 and PC2 capture 73.62% and 15.14% of the data variance, respectively. b) Line graph showing patterns of immobility with mean freeze duration activity for *Panx2^+/+^* (red) and *Panx2^Δ11^* (blue) over alternating 20-minute periods of light-ON and light-OFF conditions. c) Line graph showing the mean average activity. Mean activity, with error bars showing SD for d) light-ON and e) light-OFF conditions, ****P-value<0.0001. f) Line graph showing the mean average activity for *Panx2^+/+^* (red) and *Panx2^Δ11^* (blue) during an initial light-OFF period, followed by 20-minute increments of increasing light intensities. Mean activity, with error bars showing SD during g) 10% light intensity, h) 20% light intensity and i) 30% light intensity. ****P-value<0.0001. Data shown for n=24 larvae for each genotype.

A modified VMR assay was performed next where the intensity of light was gradually increased; locomotor activity was measured at 10% (400 lux), 20% (800 lux) and 30% (1200 lux) light intensity. With the onset of light, *Panx2^+/+^* larvae exhibited a progressive increase in average activity duration. In contrast, *Panx2^Δ11^*larvae activity was less variable as the light stimulus intensified (**Fig. 6f**). The variance analysis revealed significant differences (P-values <0.0001) in activity between *Panx2^+/+^* and *Panx2^Δ11^* larvae across all levels of luminosity (**Fig. 6g,h,i**). Altogether, the differential responses demonstrated that *Panx2^Δ11^* larvae can sense changes in light but their observed motor response to luminance is altered.

### Panx2 ablation alters the optomotor response of larvae

Optomotor response (OMR) assays evaluated the visual perception of *Panx2^Δ11^* larvae. The assay enabled us to test an innate behavioral response that many animals use to stabilize themselves relative to a visual environmental stimulus ^24^. Both *Panx2^+/+^* and *Panx2^Δ11^*larvae responded to the presented visual stimuli, composed of moving black and white stripes. Previously optimized parameters ^25^ for three spatial frequencies (SF: 64, 128, and 256 pixels/cycle), two velocities (72 and 144 pixels/sec), and two contrast levels (10% and 100%) were tested. The line graphs in **Fig. 7** summarize the percentage of positive response (PPR); corresponding to the number of larvae that showed the expected response by swimming in the direction of the stimulus (n=16 for genotype, for each parameter). Altogether, *Panx2^Δ11^*larvae demonstrated lower PPR and hence lower visual acuity in comparison to the wildtype group for all permutations of SF, speed, and contrast.

**Figure 7:**
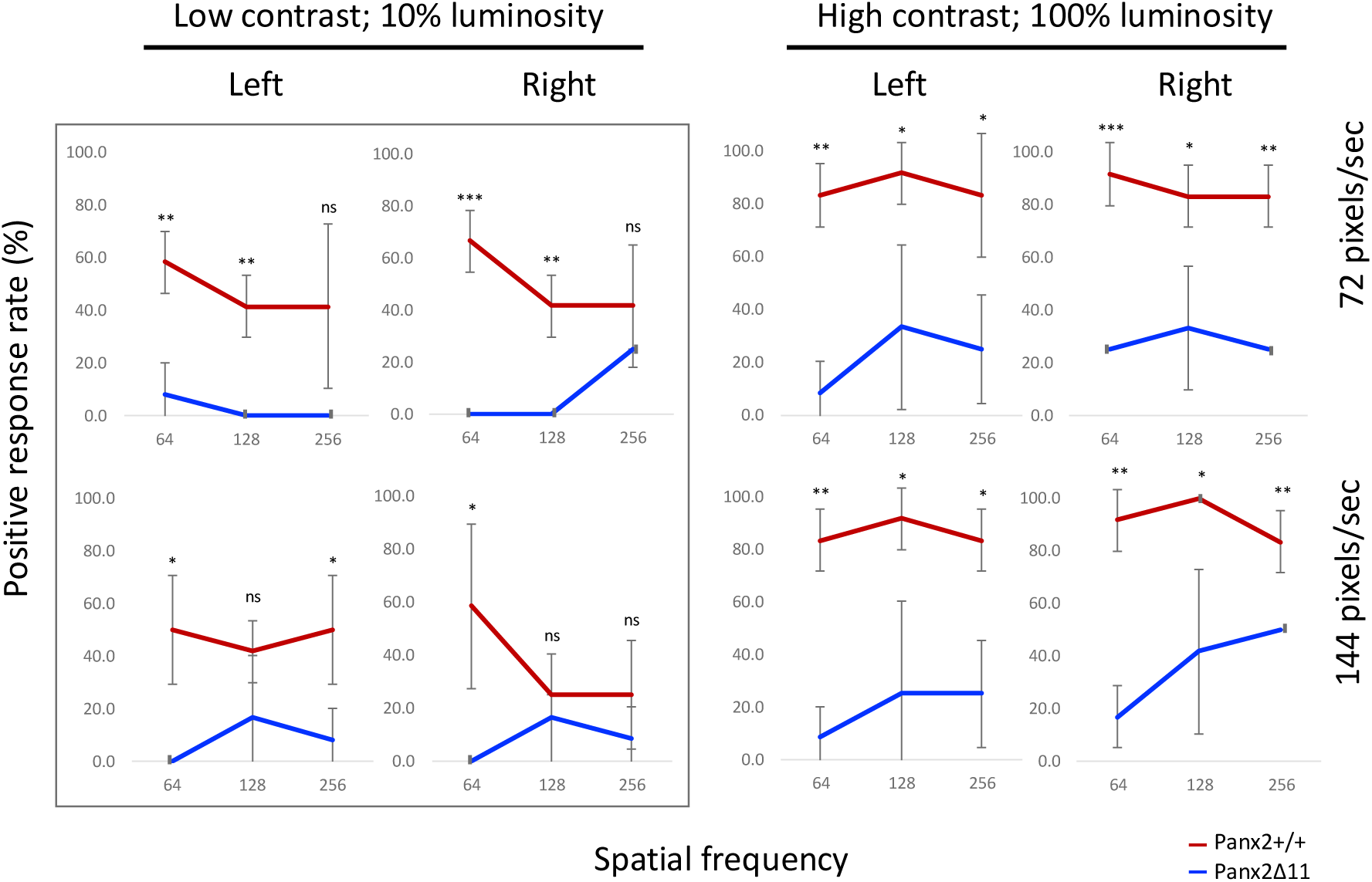
Optomotor response (OMR) impaired in *Panx2^Δ11^* larvae. OMR was used to assess optic flow processing in 6 dpf *Panx2^+/+^* and *Panx2^Δ11^*larvae, at two light intensities (10% and 100%). Both groups were shown visual stimulus of moving stripes at three spatial frequencies (64, 128, and 256), which were presented at 72 pixels/sec and 144 pixels/sec. The line graphs show the percentage of larvae that indicated the expected positive response, from n=16 for each genotype. *Panx2^Δ11^* larvae had significantly lower visual acuity when compared to *Panx2^+/+^*. ***P-value<0.001, **P-value<0.01, *P-value<0.05, ns = not a significant difference.

Under low contrast settings (10%), *Panx2^Δ11^* larvae showed an average PPR of 6.94 in comparison to 45.15 for *Panx2^+/+^*. The variance was most prevalent for the rightward direction at a speed of 72 pixels/sec and SF of 64, where *Panx2^Δ11^*larvae did not respond while PPR for TL was 66.7 (P-value 0.00061, n=16 for each genotype). Similarly, when the stimulus was presented in the leftward direction at the same speed and SF, *Panx2^Δ11^*larvae had a PPR of 8.3 in comparison to a much higher value of 58.3 for *Panx2^+/+^* (P-value 0.006, n=16 for each genotype). No statistically significant difference was observed between the two groups at the SF of 256 at 72 pixels/sec; and with the SF of 128 at 144 pixels/sec, regardless of directionality.

In the high contrast setting (100%), *Panx2^Δ11^*larvae showed an average PPR of 26.38. In this luminous environment, a statistically significant difference was observed between *Panx2^+/+^*and the knockout larvae at all combinations of SF and speed. Furthermore, peak response deficiency was observed for *Panx2^Δ11^* larvae at a SF of 64, reflecting the trend observed under low contrast conditions at the same SF. While there was an increase in PPR for *Panx2^+/+^* at high contrast, when the edges of the moving stripes were well defined, there was still a considerable decrease in OMR response with the loss of Panx2. The observed decline in visual acuity of *Panx2^Δ11^* larvae was found to be independent of direction (P-values: see Table 1).

**Table 1:** Comparison of OMR positive response percentages between *Panx2^+/+^* and *Panx2^Δ11^* larvae.

|  |  | Leftward |  |  | Rightward |  |  |
| --- | --- | --- | --- | --- | --- | --- | --- |
|  |  | Spatial frequency (pixels/cycle) |  |  |  |  |  |
| Contrast | Speed (pixels/sec) | 64 | 128 | 256 | 64 | 128 | 256 |
| Low (10%) | 72 | 0.006 | 0.003 | 0.08 | 0.00061 | 0.004 | 0.29 |
|  | 144 | 0.013 | 0.176 | 0.037 | 0.03 | 0.575 | 0.287 |
| High (100%) | 72 | 0.001 | 0.04 | 0.03 | 0.00061 | 0.03 | 0.001 |
|  | 144 | 0.0014 | 0.036 | 0.012 | 0.0014 | 0.031 | 0.008 |

### Panx2 loss leads to lens malformation consistent with myopia

Whole-eye Optical Coherence Tomography (OCT) images of adult, one-year-old, *Panx2^+/+^* and *Panx2^Δ11^* zebrafish were acquired using a custom-developed 1310nm Spectral Domain OCT (SD-OCT) system. An analysis of each eye was performed from a microstructural perspective and using geometric parameters, including axial length, lens diameter, retinal radius, and corneal thickness, as illustrated in **Fig. 8a**. Detailed examinations of whole-eye images revealed disruptions to the lens epithelium in *Panx2^Δ11^*fish. These disruptions manifested as protruding rings within the lens, unlike the smooth, well-organized epithelium observed in *Panx2^+/+^*fish (**Fig. 8b,c,d**). We speculated that the defects could impair the lens’s light-focusing ability by disrupting the normal light pathway.

**Figure 8:**
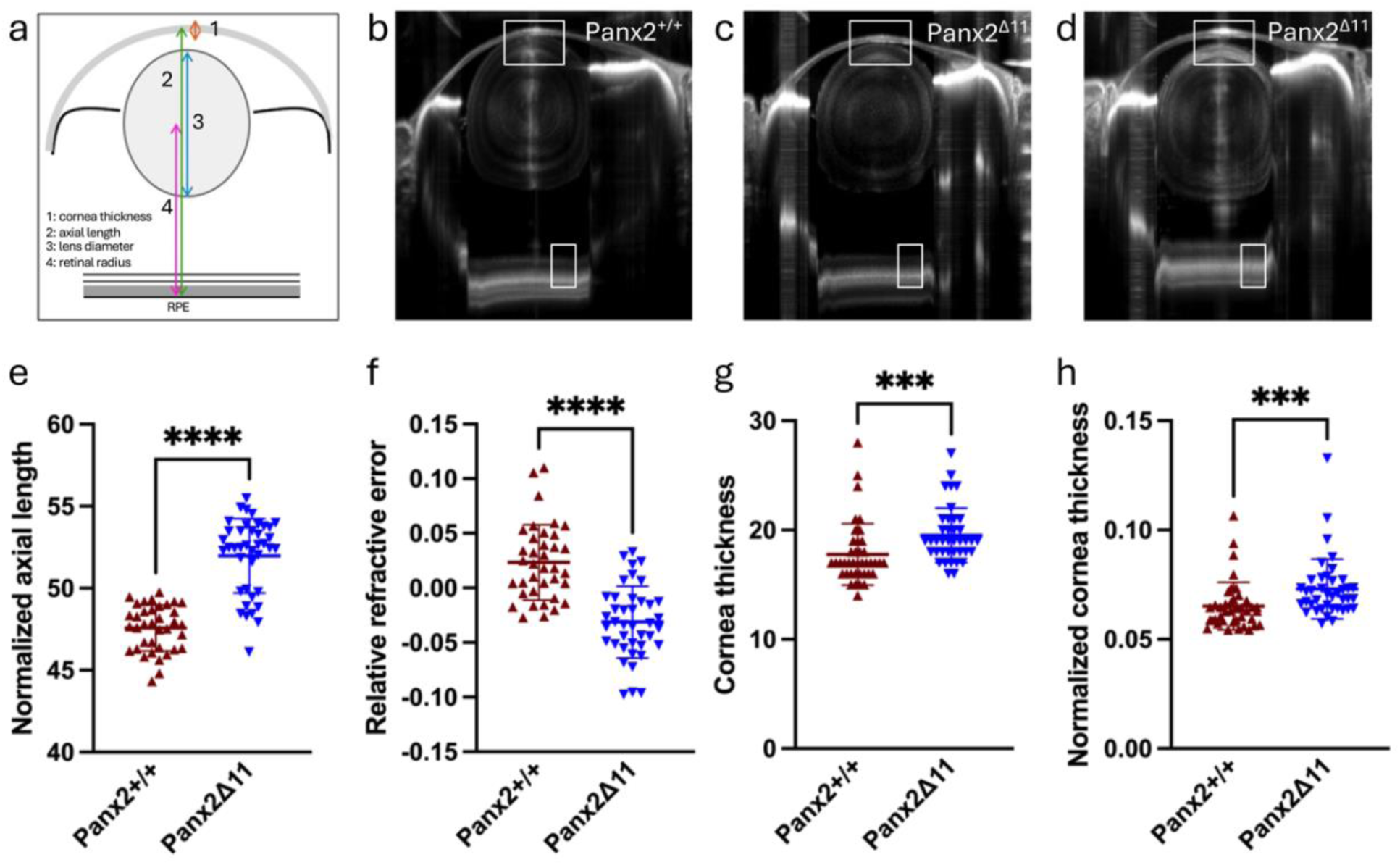
Loss of Panx2 leads to lens defects. Eyes of adult fish were imaged using a SD-OCT system. a) Schematic definition of four geometric parameters used for quantification of high-resolution OCT Whole-eye images. Representative images of *Panx2^+/+^* (b) and *Panx2^Δ11^* (c,d), showing disruptions in lens epithelium. Scale bars = 500 μm. Bar plots showing spread of quantified geometric parameters for the axial length normalized to body length (e), relative refractive error (f), corneal thickness (g), and corneal thickness normalized to lens diameter (h), collectively showing significant differences between the genotypes. Sample sizes for *Panx2^+/+^* and *Panx2^Δ11^* were 38 and 40 eyes from 19 and 20 fish, respectively. Significance: ***P-value <0.001 and ****P-value <0.0001.

To account for individual variations in body size, we normalized geometric data to body length as previous described ^26,27^. Our analysis revealed a significantly longer mean axial length in *Panx2^Δ11^*fish compared to wild-type controls (*Panx2^+/+^* = 47.57 ± 1.39 µm/mm; *Panx2^Δ11^* = 51.97 ± 2.26 µm/mm; P-value < 0.0001; Fig. 8e). To further investigate potential refractive anomalies, we calculated the Relative Refractive Error (RRE) using high-resolution whole-eye OCT images. RRE is a widely used standardized metric calculated as 1- (retinal radius/F), where F is an idealized focal length equal to lens radius × 2.324. Examination of calculated RREs revealed that Panx2 ablation led to a decrease in RRE values (*Panx2^+/+^* = 0.02 ± 0.03; *Panx2^Δ11^* = -0.03 ± 0.03; P-value <0.0001; **Fig. 8f**). Eyes with a distance from the lens center to the retinal pigment epithelium (RPE) greater than the expected retinal radius exhibit negative RRE values, indicating myopic shifts. Conversely, eyes with a shorter distance (RPE closer to the lens) have positive RRE values, signifying hyperopic shifts ^26^. The observed statistically significant reduction of RRE suggested myopia-like abnormalities in *Panx2^Δ11^* fish. Finally, we detected an increase in corneal thickness in *Panx2^Δ11^* fish, when normalized to lens diameter (*Panx2^+/+^* = 0.07 ± 0.01 µm/mm, n=38 eyes; *Panx2^Δ11^*= 0.08 ± 0.01 µm/mm, n=40 eyes; P-value <0.001; **Fig. 8g,h**). For the above-mentioned parameters, we examined a total of 38 eyes from 19 *Panx2^+/+^* control and 40 eyes from 20 *Panx2^Δ11^* age-matched adult fish. Altogether, our analysis demonstrated lens abnormalities in *Panx2^Δ11^* fish, which suggested that Panx2 loss-of-function may contribute to myopic vision deficits in adult zebrafish.

## Discussion

This study demonstrates the roles of Panx2 for visual function in zebrafish. We provide three lines of evidence. We show that Panx2 mRNA is elevated within visual processing regions of the brain; this is further supported by RNA-seq analysis that shows the differential expression of genes with biological functions in the sensory perception of light. The testing of behavior in response to different light stimuli demonstrates that Panx2-deficient larvae exhibit altered locomotor behavior in response to illumination and impaired optomotor responses, indicating defects in visual perception and acuity. Finally, in adult zebrafish, loss of Panx2 is correlated with changes to the geometry and refractive properties of the eyes.

The novel results advance our previous work on pannexin functions in the visual system. We had shown previously functions of two other pannexin family members. Panx1a is predominantly expressed in the outer retina and functions in light decrement detection ^28,29^. Our work on Panx1b revealed that the isoform is localized to the inner retina and ganglion cell layer ^25,30^. We also showed that Panx1b is expressed in the end-feet of Muller glial cells, which was the first evidence for a unique cell type-specific expression, - neuronal versus glial -unique to fish ^25^. Behavioral analyses have revealed distinct phenotypes for Panx1a and Panx1b knockout zebrafish. Panx1a KOs exhibited altered visual motor responses (VMR), characterized by a general reduction in activity and specific deficits in dark-phase locomotion ^29^. Panx1b knock outs, on the other hand, showed impairments in optic flow direction-selectivity, particularly at high spatiotemporal frequencies and low contrast surroundings ^25^. It was also demonstrated that Panx1a and Panx1b function in the outer retina as the slow negative feedback signal from HCs to cones to generate the center/surround organization of bipolar cell receptive fields ^31^. Here, we show that Panx2 is expressed in the inner and outer retinal layers, with prominent expression in the photoreceptor inner segment. Our data aligns with a previous report in the murine model, which showed that Panx2 is expressed in the photoreceptor inner segment and outer plexiform layer of the mouse retina ^7^. Panx2 KO larvae exhibit a pronounced reduction in locomotor activity during light phases and an increase in locomotor activity during dark phases, a pattern that is markedly different from the VMR phenotypes observed in Panx1a/b KOs ^25,29^.

Furthermore, regardless of direction, the OMR response in Panx2 KOs is severely impaired during low and high contrast conditions. Thus, our findings for Panx2 KO larvae suggest a severe and distinct role in retinal function compared to Panx1a/b. Consistent with the behavioral data, RNA-seq analysis identified a marked decrease in the expression of genes involved in detecting light stimuli in Panx2 knockouts. We speculate that Panx2 plays a critical role in retinal signaling that is not redundant with Panx1a or Panx1b. Their specific functions may likely be complementary or synergistic, contributing to different aspects of visual processing.

The observed expression patterns and widespread changes in gene expression and behavior highlight the pivotal role of Panx2 in retinal physiology. Its strategic localization at ER-mitochondria contacts (MAMs) and its abundance in the neuropil, a critical region for synaptic transmission, suggest that the neuronal deficits in Panx2 knockout fish could stem from bioenergetics and synaptic function disruptions. Given their highly compartmentalized nature, neurons rely on precise calcium control that is essential for their function. The endoplasmic reticulum (ER) and mitochondria are critical for this regulation ^32^. Mitochondria, in addition to providing energy to postsynaptic spines, also help maintain calcium homeostasis ^33^. We reason that the loss of Panx2 function in the inner segments of photoreceptors affects the biosynthetic machinery essential for photoreceptor function. Inner segments that are rich in mitochondria, can interface with MAMs, and consequently serve as conduits for calcium ion flux and other signaling events between organelles. Studies indicate that dysfunction in the inner segment’s ability to produce energy can lead to vision loss and degenerative diseases like retinitis pigmentosa and age-related macular degeneration ^34^.

Additionally, disruptions in the inner segment’s metabolic processes are associated with stress responses that may lead to photoreceptor cell death, impacting vision over time ^35^. As revealed by recent imaging techniques, the intricate network of ER-mitochondria connections underscores the importance of MAMs in regulating calcium concentrations and energy dynamics within neurons ^17^. We know that the metabolic demands of synaptic transmission are substantial, requiring a constant supply of ATP ^36^. It is possible then that Panx2-mediated processes within the ER-mitochondrial axis influence energy production or utilization, leading to deficits in neuronal function. Additionally, the enriched expression of Panx2 in the neuropil suggests its involvement in regulating synaptic transmission, potentially through its role in calcium signaling or mitochondrial dynamics, which are known to influence the efficiency of energy production. Studies have shown that ER releases calcium in dendrites upon synaptic stimulation, which is subsequently buffered by mitochondria.

Moreover, the tethering of mitochondria to the ER might influence the duration of ER residence in spines, potentially affecting synaptic plasticity ^37^. Computational modeling has further emphasized the role of MAMs in regulating calcium and ATP dynamics within neurons, highlighting the importance of MAMs communication in shaping synaptic signaling ^32^. Interestingly, prior overexpression studies revealed that both paralogs of Panx2, Panx1 and Panx3, can form calcium-permeable ER channels ^38,39^. These possibilities allude to the role of Panx2 in supporting retinal function and indicate that its influence likely extends beyond the visual signaling pathway.

The expression of Panx1 and Panx2 has been previously identified in the adult murine lens epithelium ^40^. In adult zebrafish, the ablation of Panx2 ablation resulted in lens malformations consistent with myopia. RNA-seq data collaborated this finding, indicating differential expression of genes associated with the structural components of the eye lens early in life. The downregulation of genes for lens-specific proteins, including alpha- and beta/gamma-crystallins and fish-specific M-subfamily gamma-crystallins, was particularly significant. These genes are essential for maintaining lens structure, and mutations in them can lead to cataracts ^41,42^. Our study suggests that Panx2 is critical for cellular energy homeostasis. The loss of Panx2 could disrupt the energy supply to lens cells, hindering their growth and differentiation and potentially resulting in a myopic phenotype. The cellular stress induced by metabolic disruptions may contribute to the development of myopia-related ocular changes, such as altered lens curvature or axial elongation. These potential implications of our research open new avenues for further investigation and underscore the multifaceted role of Panx2 in retinal physiology.

Beyond its impact on vision, transcriptome profiling also suggested that Panx2 is a novel regulator of immune responses, with immune activation potentially exacerbating retinal and lens degeneration in aged animals. While we have yet to prioritize investigating this result, Panx2 plays multifaceted roles in maintaining visual acuity, regulating ocular morphology, and moderating neuro-immune interactions crucial for retinal and lens development and health. In this regard this study’s implications extend to understanding the neuro-immune axis in vision and suggest that Panx2 could be a therapeutic target for conditions involving retinal degeneration, immune dysregulation, cataracts, or myopia.

## Materials and Methods

### Zebrafish lines

Zebrafish (*Danio Rerio*) of strain Tubingen long fin (TL) were maintained in groups with mixed sex in a recirculation system (Aquaneering Inc., San Diego, CA) at 28°C on a 14hr light/10hr dark cycle. All animal work was performed at York University’s zebrafish vivarium and in an S2 biosafety laboratory following the Canadian Council for Animal Care guidelines after approval of the study protocol by the York University Animal Care Committee (GZ#2019-7-R2).

### Generating *Panx2^Δ11^* zebrafish

Potential TALENs target sites were identified using Mojo Hand software (http://talendesign.org) ^11,12^. TALENs targeted exon 1 of the *panx2* gene (NM_001256641). The TALEN constructs were synthesized in Dr. Stephen Ekker’s lab (Mayo Clinic Cancer Center, Rochester, MN). TALEN assemblies of the repeat-variable di-residues (RVD-containing repeats) were conducted using the Golden Gate approach ^43^. pT3TS-GoldyTALEN expression vectors ^44,45^. Plasmids were linearized with the SacI restriction endonuclease (ThermoFisher Scientific, Canada) for 15 min at 37°C and used as templates for *in vitro* transcription.

Capped cRNAs were synthesized from TALEN pairs mixed 1:1 using the mMESSAGE mMACHINE T3 Transcription kit (Life Technologies, Canada) and purified using the Oligotex mRNA Mini Kit (Qiagen Inc., Toronto, Canada). TALEN cRNAs were diluted in DNase/RNase-free water (Life Technologies) to the final concentration of 1 µg/μL and stored at −80°C before microinjection.

One-cell stage zebrafish embryos were microinjected with TALEN cRNAs pair at a dose of 25 pg/nl. Genomic DNA (gDNA) was extracted from injected embryos at 4 dpf to examine the TALEN mutagenesis efficiency. Individual larvae were incubated in 100 mM NaOH at 95℃ for 15 min. After cooling to room temperature, one-tenth of the 1 M Tris (pH8.0) volume was added to the extracts to neutralize the NaOH. Finally, 1 volume TE buffer pH 8.0 was added, and gDNAs were stored at -20℃. gPCR was used as a screen to detect small indel mutations by BamHI restriction enzyme (RE) digests. PCR primers for genotyping: 5’-CGAATGCAGAATATCCTCGAGCAG-3’; reverse, 5’-GTGACGACCCGGTCAAAGG-3’. Indel mutations were confirmed by sequencing (Eurofins Genomics LLC, KY, USA) of gel-purified PCR products cloned into the pJet1.2 cloning vector (Life Technologies).

Adult mosaic zebrafish (F0) were anesthetized in pH-buffered ethyl 3-aminobenzoate methanesulfonate solution (0.2 mg/ml; MS-222, Sigma-Aldrich). A section of the caudal fin was removed using dissecting scissors (WPI Inc., FL, USA) and placed into 1.5 ml collecting tubes. The fin gDNA was isolated and screened for indel mutations as described ^44^. Adult F0 zebrafish were out-crossed to wild-type (WT) TL zebrafish and their F1 offspring were analyzed by PCR and *BamHI* restriction digestions to verify germline transmission of mutations. Heterozygous *Panx2^+/-^*F1 mutants were in-crossed to establish homozygous F2 mutants *Panx2^Δ11^*. All experiments described were performed with progenies of F3 or alter generations. Age matched siblings served as controls.

### NGS RNA-sequencing

The transcriptomes of the *Panx2^Δ11^* and sibling controls (*Panx2^+/+^*) were analyzed by RNA-seq (NGS-Facility, The Center for Applied Genomics, SickKids, Toronto, ON, Canada). The raw data represented the sequencing of three independent pools of ≈30 age-matched larvae at 6 dpf. The RNA-seq data are deposited at the NCBI - Gene Expression Omnibus (GEO) database repository (ID pending). Total RNAs were extracted using RNeasy Plus Mini Kit (Qiagen). The RNA quality was determined using a Bioanalyzer 2100 DNA High Sensitivity chip (Agilent Technologies, Mississauga, ON, Canada). The RNA library preparation was performed following the NEB NEBNext Ultra II Directional RNA Library Preparation protocol (New England Biolabs Inc., Ipswich, MA, USA). RNA libraries were loaded on a Bioanalyzer 2100 DNA High Sensitivity chip to check for size, quantified by qPCR using the Kapa Library Quantification Illumina/ABI Prism Kit protocol (KAPA Biosystems, Wilmington, MA, USA). Pooled libraries were paired-end sequenced on a High Throughput Run Mode flow cell with the V4 sequencing chemistry on an Illumina HiSeq 2500 platform (Illumina, Inc., San Diego, CA) following Illumina’s recommended protocol to generate paired-end reads of 126-bases in length.

### Differential Gene Expression and Functional Enrichment Analyses

The post-sequencing processing to final read counts, normalization, and differential gene expression analysis used multiple software packages, including a two-condition differential expression analysis using the edgeR R-package, v.4.0.16 ^46,47^ and DESeq2 R package version 1.42.1 ^48^. Genotypes were incorporated into the statistical model and multiple hypothesis testing was performed. The default filter for DESeq2 used a threshold of p<0.05 and the Benjamini-Hochberg procedure to determine the false discovery rate (FDR), and the adjusted P-value (padj).

A Gene Set Enrichment Analysis (GSEA) tested whether a defined set of genes shows statistically significant, concordant differences between genotypes. The data were filtered at padj <0.05 and analyzed for enrichment using R GOseq (v1.56.0) ^23^. Other tools used to process RNA-seq data were HCOP (https://www.genenames.org/tools/hcop/) and db2db at bioDBnet (https://biodbnet.abcc.ncifcrf.gov/db/db2db.php) to convert curated human or mouse to zebrafish genes ^49,50^. Curated gene lists were generated based on GOseq and analyzed in STRING v12.0 (string-db.org) ^51^. The top-scoring categories were visualized using ggplot2 (v3.5.1) from the tidyverse package (v2.0.0) in R ^52^.

### Hybridization chain reaction RNA-fluorescence in situ hybridization (HCR RNA-FISH)

Larvae were raised under standard conditions in egg water. When embryos reached 12 hours post-fertilization (hpf), egg water was replaced with egg water containing 0.003% of 1-phenyl 2-thiourea (PTU). Fresh egg water containing 0.003% PTU was replaced every 24 hours until the larvae reached 5 dpf. At 6 dpf, the larvae were euthanized and fixed in 4% ice-cold paraformaldehyde (PFA). After 24 hours, the PFA was washed out with 1× PBS, and the samples were gradually dehydrated, permeabilized with methanol, and stored at −20°C for several days until HCR in situ labelling was performed.

Staining was performed according to the manufacturer’s protocol for whole-mount zebrafish larvae ^53^. Samples were separated into 5 larvae per well in a 24-well plate. Rehydration steps were performed by washing for 5 minutes (mins) each in 75% methanol/PBST (1× PBS + 0.1% Tween-20), 50% methanol/PBST, 25% methanol/PBST, and 5 times with 100% PBST. The samples were permeabilized with 30 µg/ml proteinase K for 45 mins at room temperature (RT), followed by post-fixation with 4% PFA for 20 mins at RT, and 5 washes in PBST for 5 mins each. The samples were prehybridized in 500 µl of probe hybridization buffer (Molecular Instruments) for 30 mins at 37°C. Hybridization was performed by adding 2 pmol of each probe set to the hybridization buffer and incubating for 16 hours at 37°C. Probe sets for *panx2* were purchased from and designed by Molecular Instruments using proprietary HCR methodology to detect and fluorescently label target RNA transcripts. To maximize targeting, the probe was designed against shared regions of known variants found on NCBI Gene and the Ensembl database. Each probe set consisted of 20 split-initiator probe pairs per target and utilized a B1 amplifier with a 546 nm fluorophore label. To remove excess probes, the samples were washed 4 times for 15 mins each with a wash buffer (Molecular Instruments) at 37°C, followed by 2 washes of 5 mins each with 5× SSCT (5× SSC + 0.1% Tween-20) at RT. Pre-amplification was performed by incubating the samples for 30 mins in an amplification buffer (Molecular Instruments) at RT. The fluorescently labelled hairpins (B2-488) were prepared by snap cooling: heating at 95°C for 90 seconds and then cooling to RT for 30 mins. The hairpin solution was prepared by adding 10 µl of the snap-cooled hairpins (3 µM stock concentration) to 500 µl of amplification buffer. The pre-amplification buffer was removed, and the samples were incubated in the hairpin solution for 16 hours at RT. Excess hairpins were washed three times with 5× SSCT for 20 mins each. Following HCR RNA-FISH, larvae were stained with DAPI (1:12,000) overnight at 4°C, followed by three 10-minute washes in 5× SSCT. The samples were then stored in 5× SSCT in the dark at 4°C until imaging. A negative control was included, where no probes were added but fluorescent hairpins were used. The larvae were raised under standard conditions, with PTU treatment, euthanized at 6 dpf, and processed for HCR in situ labeling.

### Confocal image acquisition and analysis

A z-mold or an 8-teeth mold for zebrafish larvae was 3D-printed using 2% low melting agarose on glass-bottom culture dishes (MatTek, P35G-0-10C) ^54^. The larvae were positioned in larva-shaped slots and embedded dorsal side down in 0.5% low melting agarose. Images were acquired on the Nikon A1R confocal system with 20x W objective using the 546 nm (B1 amplifier) and 468 nm lasers. Image analysis was performed using FIJI (v2.9.0).

The acquired z-stacks were mapped against the mapZebrain atlas ^20,55^ using the DAPI stain channel as a bridge (see reference brain mapping in materials and methods). Median raw fluorescence values from *Panx2* probe and control larvae were then extracted from well-defined anatomical mapZebrain atlas regions. For visualization, these regions were overlayed on panx2 stacks using FIJI. The correlating regions are displayed either as transversal or lateral sections, or as a 3D projection viewed dorsally.

### Reference brain mapping

We used the ANTs brain registration library (Avants et al. 2011) to map volumes against the mapZebrain atlas ^20,55^. We first used Fiji to manually convert each acquired confocal volume into separate nrrd files, one for the DAPI and one for the *panx2* channel. To then generate a warp transformation file across coordinate systems, we mapped DAPI volumes against the T_AVG_DAPI reference volume (downloaded from the mapZebrain atlas platform). We then applied the resulting transform to the *panx2* probe channel. We used brain region masks (downloaded from the mapZebrain platform) to slice out the fluorescence from the mapped image stacks for further analysis of the *panx2* expression levels within well-defined separate brain areas.

For all mappings, we used the following ANTs commands: antsRegistration -v 1 -d 3 --float 1 --winsorize-image-intensities [0.005, 0.995] --use-histogram- matching 0 -o <output.nrrd> --initial-moving-transform [<DAPI_reference.nrrd>,<image_DAPI_channel.nrrd>,1] -t Rigid[0.1] -m MI[<DAPI_reference.nrrd>,<image_DAPI_channel.nrrd>,1,32,Regular,0.5] -c [1000x500x250x300,1e-8,10] -s 3x2x1x0 -f 8x4x2x1 -t Affine[0.1] -m MI[<DAPI_reference.nrrd>,<image_DAPI_channel.nrrd>,1,32,Regular,0.5] -c [200x200x200x100,1e-8,10] -s 3x2x1x0 -f 8x4x2x1 -t SyN[0.1,6,0] -m CC[<DAPI_reference.nrrd>,<image_DAPI_channel.nrrd>,1,2] -c [200x100,1e-7,10] -s 4x3 -f 12x8 antsApplyTransforms -d 3 -v 0 -- float -n linear -i <image_panx2_channel.nrrd> -r <DAPI_reference.nrrd> -o <image_panx2_channel_mapped.nrrd> -t <image_1Warp.nii.gz> -t < image_0GenericAffine.mat>

### Quantitative real-time PCR (qRT-PCR) analysis

Total RNA (1 µg) was extracted and purified from pools of 30 zebrafish larvae at 6 dpf using the RNeasy Plus Mini Kit (Qiagen, Germantown, MD, United States of America) as per the manufacturer’s protocols. Fish were homogenized by bead beating in 1xTE buffer (pH 8.0). RNAs were reverse transcribed into cDNA using the iScript Reverse Transcription Supermix (Bio-Rad, Mississauga, ON, Canada) as per the manufacturer’s instructions. Gene expression was analyzed by qRT-PCR using the SsoAdvanced universal SYBR Green Supermix (Bio-Rad, Mississauga, ON, Canada) in the CFX96™ Real-Time PCR Detection System (Bio-Rad, Mississauga, ON, Canada). Thermal cycling was carried out for 39 cycles of the following: 94°C for 30 seconds, 50°C for 30 seconds, and 72°C for 1 min. The housekeeping gene, *18S*, was used to determine the quality of the samples and for normalization purposes. CT values were averaged and exported from the CFX Manager Software (Bio-Rad, Mississauga, ON, Canada). The fold-difference for the relative gene expression was calculated using the Relative Expression Software Tool VS2009 ^56^. Three technical replicates per gene were performed from three biological replicates.

Primer sequences used for qRT-PCR:

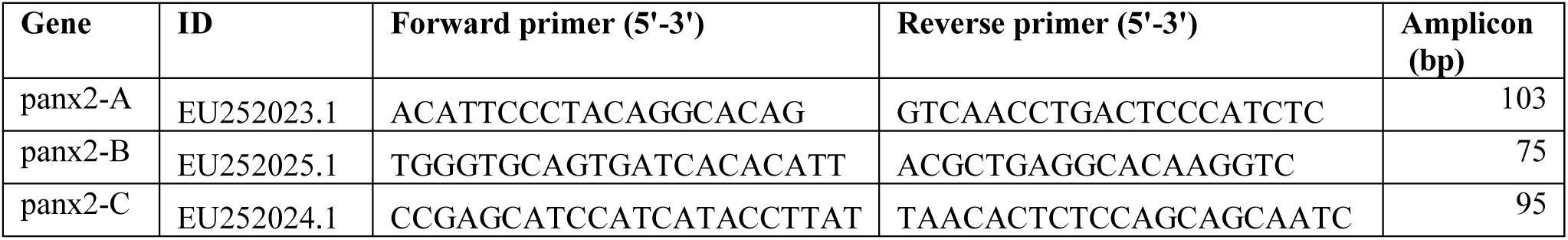

### Immunohistochemistry

Larvae (6 dpf) were humanely euthanized in MS-222 solution (0.02% w/v, Sigma-Aldrich) and fixed in 4% paraformaldehyde (PFA) in 1xPBS overnight at 4°C, followed by cryoprotection in 30% sucrose in 1xPBS. After embedding in Tissue-Tek O.C.T compound 15µm sections were cut on a cryotome (Thermofisher). Samples were washed three times for 5 mins with 1xPBS containing 0.1% Tween-20 (PBST) at RT. Unspecific binding sites were blocked with freshly prepared 5% normal goat serum (NGS, Sigma-Aldrich) in PBST for 1 hour at RT. Following blocking, samples were incubated with a custom anti-Panx2 antibody (1:200, GenScript) overnight at 4°C. A monoclonal anti-Panx2 antibody was generated against a carboxy-terminal peptide (IKAEKPKPVRRKNV), representing amino acids 460-473 of Panx2. The antibody was produced in rabbit and affinity-purified by GenScript (Nanjing GenScript Biotech Co., Ltd). Subsequent washes with PBST were carried out for 1 hour at 4℃. The Alexa 488 goat anti-rabbit secondary antibody (1:20,000 in 1% NGS PBST, catalog#A11034, Life Technologies) was applied for 1 hour at RT. After three washes with PBST and one wash with water, specimens were mounted on microscope slides using ProLong Antifade with DAPI (Thermofisher). Confocal images were collected using LSM-ZEN2 software (Zeiss LSM700 system; Carl Zeiss MicroImaging, Oberkochen, Germany) with Plan-Apochromat 20x/0.8 or Plan-Apochromat 63x/1.3 oil DIC M27 objectives. For comparison of wild type and knockout tissues settings for raw image collection were identical. Post image collection composite figures were created using Adobe Photoshop 2021.

### Zebrafish Locomotion Assays

The Zebrabox behavior system and the ZebraLab analytical suite (ViewPoint Life Technology, Lyon, France, http://www.viewpoint.fr) were used for automated extraction of behavioral outputs and video tracking. Tracking videos were recorded at 30 frames per second (fps) under infrared (for light-OFF recording) or visible light (light-ON) illumination using a Point Grey Research Dragonfly2 DR2-HIBW camera (Teledyne-FLIR, Burlington, ON, Canada). Inside the Zebrabox a lightbox provided visible and infrared light from below for recordings using 0% to 30% light intensity (visible range, 0-1200 lux). 6 dpf larvae were observed in 24 or 48-well plates maintained at 28°C throughout the experiment. All experiments were performed between 12:00 to 2 pm, as larval (6 dpf) activity was previously reported to reach a stable level by early afternoon.

### Spontaneous free-swimming assay

Larval swimming activity under constant light-ON/OFF conditions was tested using 24-well plates. Locomotor behavior was tracked for 30 mins using the Zebrabox system. For analysis of locomotion, three thresholds were defined: slow (<2mm), medium (2-20mm), and fast (>20mm). The mean total distance traveled (mm) and velocity (mm/sec) in two swim speeds *(medium and fast)* were used for statistical analysis.

### Visual Motor Response (VMR) assay

Zebrafish larvae (n=24 per genotype) were acclimatized to darkness for 2 hours in a 48-well plate. Baseline activity was recorded for 21 mins under light-OFF conditions. For the first experiment, larvae were subjected to alternating 20-minute light-ON (30%, 1200 lux) and light-OFF periods, totaling 1 hour and 41 mins. For the second experiment, a modified protocol involved incremental 10% light intensity increases every 20 mins, culminating in 30% (1200 lux) light intensity after 1 hour and 21 mins. Data acquisition for all experiments was conducted using the Quantization® mode of Zebrabox. Data collected at each time point included freeze count, freeze duration, mid count, mid duration, burst count, and burst duration. For analysis, the average duration (mean of mid and burst durations) was calculated for each larva.

### Opto Motor Response (OMR)

OMR assays were performed using a custom-built system, which follows the design described ^57^. Briefly, stimuli were presented from underneath the larvae using an ASUS P3B 800-Lumen LED portable projector (https://www.asus.com/ca-en/Projectors/P3B/). The larval movements were recorded using a USB 3.1 high-speed camera (XIMEA GmbH, Germany) equipped with a 35mm C Series Fixed Focal Length Lens (Edmund Optics Inc., USA). An 830 nm long-pass filter (Edmund Optics Inc., USA) was used to block infrared illumination coming from a source at the bottom of the test environment. The videos were recorded using the XIMEA Windows Software Package (https://www.ximea.com/support/wiki/apis/software_packages). Visual stimuli were generated with an online stimulus generator program called “Moving Grating” (available at https://michaelbach.de/sci/stim/movingGrating/index.html).

During experiments four larvae in a 3 cm dish (Thermo Scientific) were allowed to acclimatize for 5 mins before starting the video recording. The visual stimulus consisted of sequences of alternating white and black stripes generated with 64/128/256 pixels/cycle spatial frequency. The speed rate was set to 72/144 pixels/sec. The contrast was set to 10%, where the stripes appeared as white and grey, or 100%, where the stripes appeared as white and black. Stimuli were presented to larvae (n = 4, for each of the four independent experimental repeats) for 30 seconds in the left or right direction. Once the larvae oriented towards the moving stimulus and initiated a sustained swimming motion in the direction of the stimulus, it was counted as a positive response. The positive rate of response was used for analysis. This value was expressed as a percentage: number of larvae that swam in the direction of the stimulus/ total number of larvae in the dish.

### Optical coherence tomography (OCT) assay

A custom-developed spectral-domain optical coherence tomography (SD-OCT) system was built in a Michelson configuration, employing a super luminescent laser diode centered at 1,310 nm (± 75 nm at 10 dB; Exalos, Switzerland) and a 2048-pixel line scan camera spectrometer with a maximum acquisition rate of 147kHz (Wasatch Photonics; United States of America). A 50/50 fiber coupler splits the source light into the reference and sample arms. In the sample arm, the output light illuminates the sample surface after passing through a reflective beam collimator (Thorlabs; United States of America), a 2-DOF galvo mirror, and an objective lens (LSM02, Thorlabs; United States of America). The 2-DOF Galvo mirror allows for collection of reflected light from the sample while raster scanning sample surface. In the reference arm, a polarization controller, a dispersion compensation block, and a gold-coated reference mirror were installed. The back-reflected light of these two arms is subsequently merged after passing through the beam splitter and redirected to the spectrometer by the optical circulator. The formed interference pattern in the spectrometer is captured by a line scan camera. The captured signal is digitized and sent to the computer for processing. To form an A-line (i.e., depth profile of sample reflections at a given point on sample surface), the tomograms of the sample is background subtracted and mapped to k-space before applying Fourier transformation to calculate depth profile of reflectors in the z-space (physical depth space). Processes were repeated for data acquired during raster scanning of beam on sample surface to eventually form 3-D OCT volumetric images of zebrafish eye. The axial and lateral resolutions of the system in tissue were measured as 8.5 μm and 10 μm, respectively. Before imaging, age-matched adult zebrafish were humanely euthanized using MS-222. Fish were then placed in a silicon mold to orient the eye toward OCT system’s objective lens. To minimize specular reflections from sample surface, a thin layer of PBS (∼80 μm) was placed over the eye before imaging. All captured OCT images were calibrated for image pixel size in axial and lateral directions before interrogation in ImageJ software and quantification of the geometric parameters.

### Statistics and data reproducibility

Statistical analyses were performed in GraphPad Prism VS10.2.3. Results are represented as the mean ± standard deviation (SD) or standard error of the mean (SEM) for behavioural data. For molecular analysis (RT-qPCR, RNA-seq), a minimum of n ≥ 3 independent experimental replicates were generated. For behavioral testing G*power analysis determined the number of larvae. The normality and homogeneity of the data variance were determined by the “Shapiro-Wilk Test” and “Levene’s Test.” Experimental groups were compared using unpaired t-tests, Welch’s t-tests, or two-way ANOVA. A *P* value <0.05 was considered statistically significant. For all experiments, sample sizes, statistical tests, and when appropriate P-values are indicated in the figure legends.

## Acknowledgements

We thank two members of York University Zebrafish Vivarium, Janet Fleites-Medina, and Veronica Scavo for outstanding zebrafish husbandry. We also wish to thank the Center for Applied Genomics, SickKids, Toronto, ON, Canada for the RNA-seq service.

## Author contributions

Conceptualization, RS, GRZ; data analysis, RS, GSZ, FN, SS, HN, AB; investigation, all authors; writing – original draft preparation, RS, GRZ; writing – review and editing, all authors; visualization, RS, GSZ, FN, SS, AB; supervision, NT, GRZ; project administration, GRZ; funding acquisition, NT, GRZ, AB.

## Consent for publication

All authors have read and agreed to the published version of the manuscript.

## Ethics approval

All animal work was performed at York University’s zebrafish vivarium and in an S2 biosafety laboratory following the Canadian Council for Animal Care guidelines after approval of the study protocol by the York University Animal Care Committee (GZ#2019-7-R2).

## Data Availability

The RNA-seq data are deposited at the NCBI - Gene Expression Omnibus (GEO) database repository (ID pending at time of submission). Additional information necessary for the reanalysis of the data reported in this manuscript is available from the corresponding author upon request.

## Materials & Correspondence

Correspondence and material requests should be addressed to Riya Shanbhag, Georg R. Zoidl.

