## Supplementary Data for "Pannexin-2 deficiency disrupts visual pathways and leads to ocular defects in zebrafish"

Supplementary Table 1: *Sensory Perception of Light Stimulus* (GO:0050953)

| ***Gene*** | ***log2FoldChange*** | ***pvalue*** | ***padj*** | ***-log$_{10}$(padj)*** |
|---|---:|---:|---:|---:|
| crybb2 | -1.66 | 3.70E-03 | 2.31E-02 | 1.64 |
| arr3b | -1.28 | 5.61E-65 | 6.23E-62 | 61.21 |
| zgc:153846 | -1.11 | 1.17E-15 | 8.73E-14 | 13.06 |
| crygm1 | -1.04 | 1.44E-04 | 1.56E-03 | 2.81 |
| guca1g | -0.91 | 2.36E-16 | 1.93E-14 | 13.71 |
| arr3a | -0.81 | 8.12E-30 | 2.06E-27 | 26.69 |
| rgrb | -0.77 | 7.08E-14 | 4.39E-12 | 11.36 |
| guca1ab.2 | -0.75 | 3.56E-10 | 1.27E-08 | 7.9 |
| cryba1a | -0.65 | 9.68E-05 | 1.11E-03 | 2.96 |
| myo3a | -0.57 | 1.14E-13 | 6.70E-12 | 11.17 |
| grk7a | -0.57 | 1.71E-18 | 1.69E-16 | 15.77 |
| per1b | -0.56 | 7.48E-35 | 2.52E-32 | 31.6 |
| cryba1b | -0.53 | 6.70E-11 | 2.70E-09 | 8.57 |
| crygm2d18 | -0.53 | 2.80E-11 | 1.20E-09 | 8.92 |
| pde6c | -0.5 | 3.68E-12 | 1.77E-10 | 9.75 |
| crygm2d5 | -0.49 | 1.27E-06 | 2.36E-05 | 4.63 |
| crygm2d13 | -0.48 | 1.20E-10 | 4.65E-09 | 8.33 |
| crybb1l1 | -0.46 | 8.62E-09 | 2.44E-07 | 6.61 |
| pdca | -0.46 | 2.10E-04 | 2.15E-03 | 2.67 |
| opn4.1 | -0.45 | 4.78E-10 | 1.66E-08 | 7.78 |
| crygm2d20 | -0.43 | 6.54E-09 | 1.88E-07 | 6.72 |
| rhol | -0.42 | 8.43E-04 | 6.84E-03 | 2.16 |
| crybb1l2 | -0.41 | 2.89E-09 | 8.79E-08 | 7.06 |
| crygm2e | -0.41 | 1.22E-04 | 1.35E-03 | 2.87 |
| crygmx | -0.41 | 7.11E-03 | 3.84E-02 | 1.42 |
| grk1a | -0.39 | 1.91E-05 | 2.67E-04 | 3.57 |
| crygn1 | -0.39 | 5.26E-04 | 4.64E-03 | 2.33 |
| cryba1l2 | -0.38 | 1.03E-03 | 8.11E-03 | 2.09 |
| crygm2d15 | -0.38 | 7.59E-08 | 1.82E-06 | 5.74 |
| pde6ha | -0.37 | 4.92E-14 | 3.13E-12 | 11.51 |
| cryba1l1 | -0.37 | 4.55E-06 | 7.46E-05 | 4.13 |
| vit | -0.37 | 6.40E-08 | 1.56E-06 | 5.81 |
| cryba4 | -0.36 | 1.19E-05 | 1.75E-04 | 3.76 |
| crygm2d8 | -0.36 | 2.84E-09 | 8.68E-08 | 7.06 |
| rho | -0.36 | 8.30E-08 | 1.98E-06 | 5.7 |
| crygn2 | -0.35 | 4.94E-09 | 1.45E-07 | 6.84 |

| | | | | |
|---|---|---|---|---|
| crygm2d21 | -0.35 | 6.70E-08 | 1.63E-06 | 5.79 |
| rpe65a | -0.35 | 7.11E-05 | 8.44E-04 | 3.07 |
| crygm2d3 | -0.35 | 1.05E-07 | 2.43E-06 | 5.61 |
| rrh | -0.34 | 1.15E-03 | 8.90E-03 | 2.05 |
| nphp3 | -0.34 | 1.69E-05 | 2.39E-04 | 3.62 |
| crybgx | -0.34 | 7.68E-06 | 1.19E-04 | 3.93 |
| crygm2d12 | -0.33 | 6.76E-08 | 1.64E-06 | 5.78 |
| crybb1 | -0.33 | 1.48E-05 | 2.14E-04 | 3.67 |
| crygmxl2 | -0.31 | 3.48E-05 | 4.49E-04 | 3.35 |
| crygm2d10 | -0.31 | 4.10E-05 | 5.18E-04 | 3.29 |
| crygm2d14 | -0.28 | 3.52E-04 | 3.31E-03 | 2.48 |
| crygm2d7 | -0.27 | 7.29E-04 | 6.09E-03 | 2.22 |
| pdcl | -0.27 | 4.97E-06 | 8.08E-05 | 4.09 |
| unc119b | -0.27 | 3.88E-05 | 4.95E-04 | 3.31 |
| pde6gb | -0.26 | 2.35E-03 | 1.60E-02 | 1.8 |
| cryba2b | -0.24 | 7.01E-05 | 8.35E-04 | 3.08 |
| crygm2d17 | -0.24 | 4.73E-04 | 4.23E-03 | 2.37 |
| crx | -0.23 | 1.32E-03 | 9.95E-03 | 2 |
| opn1sw1 | -0.23 | 4.55E-03 | 2.72E-02 | 1.56 |
| si:dkey-57a22.15 | -0.23 | 8.73E-04 | 7.06E-03 | 2.15 |
| si:ch73-167i17.6 | -0.23 | 1.92E-03 | 1.36E-02 | 1.87 |
| adgrv1 | -0.21 | 4.81E-03 | 2.85E-02 | 1.55 |
| gucy2d | -0.21 | 5.10E-03 | 2.97E-02 | 1.53 |
| crygm2d16 | -0.21 | 4.51E-03 | 2.71E-02 | 1.57 |
| cryba2a | -0.2 | 1.99E-03 | 1.40E-02 | 1.85 |

Supplementary Table 2: *Sensory Perception* (GO:0007600)

| ***Gene*** | ***log2FoldChange*** | ***pvalue*** | ***padj*** | ***-log$_{10}$(padj)*** |
|---|---|---|---|---|
| crybb2 | -1.66 | 3.70E-03 | 2.31E-02 | 1.64 |
| arr3b | -1.28 | 5.61E-65 | 6.23E-62 | 61.21 |
| zgc:153846 | -1.11 | 1.17E-15 | 8.73E-14 | 13.06 |
| crygm1 | -1.04 | 1.44E-04 | 1.56E-03 | 2.81 |
| or6at1 | -1.03 | 1.60E-03 | 1.17E-02 | 1.93 |
| guca1g | -0.91 | 2.36E-16 | 1.93E-14 | 13.71 |
| arr3a | -0.81 | 8.12E-30 | 2.06E-27 | 26.69 |
| rgrb | -0.77 | 7.08E-14 | 4.39E-12 | 11.36 |
| guca1ab.2 | -0.75 | 3.56E-10 | 1.27E-08 | 7.9 |
| otog | -0.69 | 4.78E-12 | 2.25E-10 | 9.65 |
| gprc6a | -0.69 | 9.59E-05 | 1.10E-03 | 2.96 |
| cryba1a | -0.65 | 9.68E-05 | 1.11E-03 | 2.96 |

| | | | | |
|---|---|---|---|---|
| myo3a | -0.57 | 1.14E-13 | 6.70E-12 | 11.17 |
| grk7a | -0.57 | 1.71E-18 | 1.69E-16 | 15.77 |
| per1b | -0.56 | 7.48E-35 | 2.52E-32 | 31.6 |
| cryba1b | -0.53 | 6.70E-11 | 2.70E-09 | 8.57 |
| crygm2d18 | -0.53 | 2.80E-11 | 1.20E-09 | 8.92 |
| slc26a2 | -0.53 | 5.86E-13 | 3.14E-11 | 10.5 |
| pde6c | -0.5 | 3.68E-12 | 1.77E-10 | 9.75 |
| crygm2d5 | -0.49 | 1.27E-06 | 2.36E-05 | 4.63 |
| pigu | -0.48 | 1.01E-05 | 1.51E-04 | 3.82 |
| crygm2d13 | -0.48 | 1.20E-10 | 4.65E-09 | 8.33 |
| crybb1l1 | -0.46 | 8.62E-09 | 2.44E-07 | 6.61 |
| pdca | -0.46 | 2.10E-04 | 2.15E-03 | 2.67 |
| opn4.1 | -0.45 | 4.78E-10 | 1.66E-08 | 7.78 |
| crygm2d20 | -0.43 | 6.54E-09 | 1.88E-07 | 6.72 |
| rhol | -0.42 | 8.43E-04 | 6.84E-03 | 2.16 |
| ush1c | -0.42 | 1.99E-06 | 3.54E-05 | 4.45 |
| crybb1l2 | -0.41 | 2.89E-09 | 8.79E-08 | 7.06 |
| crygm2e | -0.41 | 1.22E-04 | 1.35E-03 | 2.87 |
| crygmx | -0.41 | 7.11E-03 | 3.84E-02 | 1.42 |
| grk1a | -0.39 | 1.91E-05 | 2.67E-04 | 3.57 |
| crygn1 | -0.39 | 5.26E-04 | 4.64E-03 | 2.33 |
| cryba1l2 | -0.38 | 1.03E-03 | 8.11E-03 | 2.09 |
| crygm2d15 | -0.38 | 7.59E-08 | 1.82E-06 | 5.74 |
| pde6ha | -0.37 | 4.92E-14 | 3.13E-12 | 11.51 |
| cryba1l1 | -0.37 | 4.55E-06 | 7.46E-05 | 4.13 |
| vit | -0.37 | 6.40E-08 | 1.56E-06 | 5.81 |
| cryba4 | -0.36 | 1.19E-05 | 1.75E-04 | 3.76 |
| crygm2d8 | -0.36 | 2.84E-09 | 8.68E-08 | 7.06 |
| rho | -0.36 | 8.30E-08 | 1.98E-06 | 5.7 |
| crygn2 | -0.35 | 4.94E-09 | 1.45E-07 | 6.84 |
| crygm2d21 | -0.35 | 6.70E-08 | 1.63E-06 | 5.79 |
| rpe65a | -0.35 | 7.11E-05 | 8.44E-04 | 3.07 |
| crygm2d3 | -0.35 | 1.05E-07 | 2.43E-06 | 5.61 |
| rrh | -0.34 | 1.15E-03 | 8.90E-03 | 2.05 |
| nphp3 | -0.34 | 1.69E-05 | 2.39E-04 | 3.62 |
| crybgx | -0.34 | 7.68E-06 | 1.19E-04 | 3.93 |
| crygm2d12 | -0.33 | 6.76E-08 | 1.64E-06 | 5.78 |
| crybb1 | -0.33 | 1.48E-05 | 2.14E-04 | 3.67 |
| crygmxl2 | -0.31 | 3.48E-05 | 4.49E-04 | 3.35 |
| crygm2d10 | -0.31 | 4.10E-05 | 5.18E-04 | 3.29 |

| | | | | |
|---|---|---|---|---|
| cldn7b | -0.31 | 3.66E-06 | 6.13E-05 | 4.21 |
| crygm2d14 | -0.28 | 3.52E-04 | 3.31E-03 | 2.48 |
| crygm2d7 | -0.27 | 7.29E-04 | 6.09E-03 | 2.22 |
| pdcl | -0.27 | 4.97E-06 | 8.08E-05 | 4.09 |
| unc119b | -0.27 | 3.88E-05 | 4.95E-04 | 3.31 |
| pde6gb | -0.26 | 2.35E-03 | 1.60E-02 | 1.8 |
| cryba2b | -0.24 | 7.01E-05 | 8.35E-04 | 3.08 |
| crygm2d17 | -0.24 | 4.73E-04 | 4.23E-03 | 2.37 |
| myo6b | -0.24 | 2.87E-03 | 1.89E-02 | 1.72 |
| crx | -0.23 | 1.32E-03 | 9.95E-03 | 2 |
| opn1sw1 | -0.23 | 4.55E-03 | 2.72E-02 | 1.56 |
| si:dkey-57a22.15 | -0.23 | 8.73E-04 | 7.06E-03 | 2.15 |
| si:ch73-167i17.6 | -0.23 | 1.92E-03 | 1.36E-02 | 1.87 |
| adgrv1 | -0.21 | 4.81E-03 | 2.85E-02 | 1.55 |
| gucy2d | -0.21 | 5.10E-03 | 2.97E-02 | 1.53 |
| crygm2d16 | -0.21 | 4.51E-03 | 2.71E-02 | 1.57 |
| cldn3c | -0.21 | 5.75E-03 | 3.26E-02 | 1.49 |
| cryba2a | -0.2 | 1.99E-03 | 1.40E-02 | 1.85 |

Supplementary Table 3: *Humoral Immune Response* (GO:0006959)

| ***Gene*** | ***log2FoldChange*** | ***pvalue*** | ***padj*** | ***-log$_{10}$(padj)*** |
|---|---|---|---|---|
| c9 | 1.01 | 4.21E-21 | 5.22E-19 | 18.28 |
| c8g | 1.3 | 8.05E-21 | 9.74E-19 | 18.01 |
| cfb | 1 | 4.64E-19 | 4.77E-17 | 16.32 |
| c8a | 1.08 | 1.87E-18 | 1.84E-16 | 15.73 |
| c7a | 1.14 | 1.06E-15 | 8.00E-14 | 13.1 |
| c6 | 0.7 | 6.24E-12 | 2.90E-10 | 9.54 |
| tfa | 0.65 | 2.29E-11 | 9.94E-10 | 9 |
| c7b | 0.64 | 2.30E-09 | 7.13E-08 | 7.15 |
| bfb | 0.89 | 4.19E-08 | 1.06E-06 | 5.98 |
| si:dkey-22f5.9 | 1.72 | 6.16E-07 | 1.22E-05 | 4.91 |
| cxcl18b | 0.7 | 4.43E-06 | 7.30E-05 | 4.14 |
| si:ch1073-280e3.1 | 1.03 | 3.15E-05 | 4.11E-04 | 3.39 |
| si:ch73-359m17.2 | 0.59 | 1.28E-04 | 1.41E-03 | 2.85 |
| c4b | 0.68 | 4.06E-04 | 3.71E-03 | 2.43 |
| c4 | 0.37 | 5.79E-04 | 5.01E-03 | 2.3 |
| cxcl8b.3 | 0.6 | 8.28E-04 | 6.75E-03 | 2.17 |
| c8b | 0.38 | 1.89E-03 | 1.34E-02 | 1.87 |

Supplementary Table 4: *Structural Component of the Lens* (GO:0005212)

| ***Gene*** | ***log2FoldChange*** | ***pvalue*** | ***padj*** | ***-log$_{10}$(padj)*** |
|---|---|---|---|---|
| crybb2 | -1.66 | 3.70E-03 | 2.31E-02 | 1.64 |
| zgc:153846 | -1.11 | 1.17E-15 | 8.73E-14 | 13.06 |
| crygm1 | -1.04 | 1.44E-04 | 1.56E-03 | 2.81 |
| cryba1a | -0.65 | 9.68E-05 | 1.11E-03 | 2.96 |
| cryba1b | -0.53 | 6.70E-11 | 2.70E-09 | 8.57 |
| crygm2d18 | -0.53 | 2.80E-11 | 1.20E-09 | 8.92 |
| lim2.1 | -0.5 | 1.71E-07 | 3.79E-06 | 5.42 |
| crygm2d5 | -0.49 | 1.27E-06 | 2.36E-05 | 4.63 |
| crygm2d13 | -0.48 | 1.20E-10 | 4.65E-09 | 8.33 |
| crybb1l1 | -0.46 | 8.62E-09 | 2.44E-07 | 6.61 |
| lim2.3 | -0.44 | 1.06E-05 | 1.59E-04 | 3.8 |
| crygm2d20 | -0.43 | 6.54E-09 | 1.88E-07 | 6.72 |
| crybb1l2 | -0.41 | 2.89E-09 | 8.79E-08 | 7.06 |
| crygm2e | -0.41 | 1.22E-04 | 1.35E-03 | 2.87 |
| crygmx | -0.41 | 7.11E-03 | 3.84E-02 | 1.42 |
| crygn1 | -0.39 | 5.26E-04 | 4.64E-03 | 2.33 |
| cryba1l2 | -0.38 | 1.03E-03 | 8.11E-03 | 2.09 |
| crygm2d15 | -0.38 | 7.59E-08 | 1.82E-06 | 5.74 |
| cryba1l1 | -0.37 | 4.55E-06 | 7.46E-05 | 4.13 |
| cryba4 | -0.36 | 1.19E-05 | 1.75E-04 | 3.76 |
| crygm2d8 | -0.36 | 2.84E-09 | 8.68E-08 | 7.06 |
| lim2.4 | -0.36 | 4.34E-05 | 5.45E-04 | 3.26 |
| crygn2 | -0.35 | 4.94E-09 | 1.45E-07 | 6.84 |
| crygm2d21 | -0.35 | 6.70E-08 | 1.63E-06 | 5.79 |
| crygm2d3 | -0.35 | 1.05E-07 | 2.43E-06 | 5.61 |
| crybgx | -0.34 | 7.68E-06 | 1.19E-04 | 3.93 |
| crygm2d12 | -0.33 | 6.76E-08 | 1.64E-06 | 5.78 |
| crybb1 | -0.33 | 1.48E-05 | 2.14E-04 | 3.67 |
| crygmxl2 | -0.31 | 3.48E-05 | 4.49E-04 | 3.35 |
| crygm2d10 | -0.31 | 4.10E-05 | 5.18E-04 | 3.29 |
| lim2.5 | -0.28 | 2.46E-03 | 1.66E-02 | 1.78 |
| crygm2d14 | -0.28 | 3.52E-04 | 3.31E-03 | 2.48 |
| crygm2d7 | -0.27 | 7.29E-04 | 6.09E-03 | 2.22 |
| cryba2b | -0.24 | 7.01E-05 | 8.35E-04 | 3.08 |
| crygm2d17 | -0.24 | 4.73E-04 | 4.23E-03 | 2.37 |
| si:dkey-57a22.15 | -0.23 | 8.73E-04 | 7.06E-03 | 2.15 |
| crygm2d16 | -0.21 | 4.51E-03 | 2.71E-02 | 1.57 |
| cryba2a | -0.2 | 1.99E-03 | 1.40E-02 | 1.85 |

Supplementary Table 5: OCT measurements for one-year-old fish

| | ***Panx2$^{+/+}$*** | | ***Panx2$^{\Delta 11}$*** | | |
|---|---|---|---|---|---|
| | ***mean*** | ***std dev.*** | ***mean*** | ***std dev.*** | ***pvalue*** |
| ***Normalized axial length (µm/mm)*** | 47.57 | 1.40 | 51.97 | 2.26 | <0.0001 |
| ***RRE*** | 0.023 | 0.03 | -0.031 | 0.03 | <0.0001 |
| ***Corneal thickness (µm)*** | 66.82 | 10.58 | 74.72 | 12.69 | 0.0002 |
| ***Corneal thickness/ lens diameter (µm/µm)*** | 0.074 | 0.01 | 0.082 | 0.02 | <0.0001 |
